# Increased biosynthesis of acetyl-CoA in the yeast *Saccharomyces cerevisiae* by overexpression of a deregulated pantothenate kinase gene

**DOI:** 10.1101/2021.05.25.445608

**Authors:** Judith Olzhausen, Mathias Grigat, Larissa Seifert, Tom Ulbricht, Hans-Joachim Schüller

## Abstract

Coenzyme A (CoA) and its derivatives such as acetyl-CoA are essential metabolites for several biosynthetic reactions. In the yeast *S. cerevisiae*, five enzymes (encoded by essential genes *CAB1-CAB5*; <u>c</u>oenzyme <u>A</u> <u>b</u>iosynthesis) are required to perform CoA biosynthesis from pantothenate, cysteine and ATP. Similar to enzymes from other eukaryotes, yeast pantothenate kinase (PanK, encoded by *CAB1*) turned out to be inhibited by acetyl-CoA. By genetic selection of intragenic suppressors of a temperature-sensitive *cab1* mutant combined with rationale mutagenesis of the presumed acetyl-CoA binding site within PanK, we were able to identify the variant *CAB1* W331R, encoding a hyperactive PanK completely insensitive to inhibition by acetyl-CoA. Using a versatile gene integration cassette containing the *TPI1* promoter, we constructed strains overexpressing *CAB1* W331R in combination with additional genes of CoA biosynthesis (*CAB2, CAB3, HAL3, CAB4* and *CAB5*). In these strains, the level of CoA nucleotides was 15-fold increased, compared to a reference strain without additional *CAB* genes. Overexpression of wild-type *CAB1* instead of *CAB1* W331R turned out as substantially less effective (4-fold increase of CoA nucleotides). Supplementation of overproducing strains with additional pantothenate could further elevate the level of CoA (2.3-fold). Minor increases were observed after overexpression of *FEN2* (encoding a pantothenate permease) and deletion of *PCD1* (CoA-specific phosphatase). We conclude that the strategy described in this work may improve the efficiency of biotechnological applications depending on acetyl-CoA.

**Key points:**

- A gene encoding a hyperactive yeast pantothenate kinase (PanK) was constructed.
- Overexpression of CoA biosynthetic genes elevated CoA nucleotides 15-fold.
- Supplementation with pantothenate further increased the level of CoA nucleotides.

## Introduction

Coenzyme A (CoA) is an indispensable and ubiquitous metabolic cofactor of numerous enzymes requiring activated carboxylic acids as CoA thioesters. Especially acetyl-CoA is needed as the energy source of the citric acid cycle and as a building block for the biosynthesis of fatty acids, sterols and several other compounds. Acetyl-CoA is also of fundamental importance for protein acetylation with special emphasis on histone acetylation as a mechanism for regulating chromatin accessibility in the course of gene activation (summarized by Galdieri et al. 2014). While higher organisms synthesize cytoplasmic acetyl-CoA by cleavage of citrate (exported from mitochondria, catalyzed by the ATP-citrate lyase ACL), *Saccharomyces* yeasts use acetyl-CoA synthetase for ATP-dependent activation of acetate (constitutive nucleocytosolic isoenzyme Acs2). Acetyl-CoA at the junction of anabolic and catabolic metabolism must be considered as a metabolite whose concentration is indicative of the cellular nutritional and energetic status (reviewed by Pietrocola et al. 2015). For yeast and cultured mammalian cells, fluctuation of acetyl-CoA concentration and its intracellular distribution has been proposed as an indicator of “fed” or “fasted” states with a decisive influence on fundamental aspects of cellular physiology (e. g. entry into a new cell division cycle; Shi and Tu 2013; 2015). Under conditions of oxidative stress, CoA can also trigger a regulatory response by covalent modification of proteins such as peroxiredoxins at cysteine residues (CoAlation; Baković et al. 2019). In biotechnology, acetyl-CoA is an essential compound for improving several strategies of metabolic engineering which may enable microbial biosynthesis of biofuels (n-butanol and farnesene; Schadeweg and Boles 2016; Meadows et al. 2016; Tippmann et al. 2016) as well as pharmaceutical compounds (hydrocortisone, polyketide antibiotics, cannabinoids and the antimalarial drug artemisinin; Szczebara et al. 2003; Paddon et al. 2013; Luo et al. 2019). The importance of malonyl-CoA for pathways related to biosynthesis of fatty acids and polyketides has been recently reviewed (Milke and Marienhagen 2020).

Comparative genomic studies have clearly shown that the biosynthesis of CoA utilizes five universal reactions, using pantothenate, cysteine and ATP as its substrates (reviewed by Leonardi et al. 2005b; Spry et al. 2008; Theodoulou et al. 2014). While vertebrates depend on the uptake of exogenous pantothenate (vitamin B_5_) as a precursor, most bacteria, yeasts, fungi and plants are able to synthesize pantothenate *de novo*, using the carbon backbone of amino acids. In the yeast *Saccharomyces cerevisiae*, uptake of pantothenate by the high-affinity plasma membrane H^+^-symporter Fen2 has been described (Stolz and Sauer 1999). Although some strains of *S. cerevisiae* may be auxotrophic for pantothenate, others are competent for *de novo* biosynthesis, using polyamines such as spermine as an unusual source of β-alanine which is required to form pantothenate from pantoate (White et al. 2001).

The initial and presumably rate-limiting step of CoA biosynthesis is ATP-dependent phosphorylation of pantothenate by a pantothenate kinase (PanK). While bacterial PanK enzymes are strongly inhibited by unacylated CoA (*coaA* gene product from *E. coli*; Vallari et al. 1987), eukaryotic pantothenate kinases (which are poorly, if at all, related to bacterial enzymes) are sensitive against inhibition by acetyl-CoA (first shown for the PanK of the fungus *Aspergillus nidulans*: Calder et al. 1999). A temperature-sensitive mutant of *S. cerevisiae* initially isolated because its fatty acid synthase (FAS) was pantetheine-free later turned out as defective for pantothenate kinase (G351S missense mutation). The corresponding wild-type gene *CAB1* (<u>c</u>oenzyme <u>A</u> <u>b</u>iosynthesis) exhibits substantial similarity to the enzyme of *A. nidulans* (43.8% identity) and is expressed at a low level. Although transcription of *CAB1* was not substantially affected by the carbon source or by availability of amino acids (Olzhausen et al. 2009), its upstream region contains two sequence motifs reminiscent of the sterol-response element (SRE) which is bound by activators Upc2 and Ecm22 (Brohée et al. 2011). Thus, it should be possible to strongly elevate yeast PanK activity by expression of *CAB1* using a heterologous promoter. While lower eukaryotes contain a single PanK gene copy, four *PANK* genes with different tissue-specific patterns of transcription could be identified in mammals, encoding enzymes with distinct cellular localization and sensitivity against CoA and acetyl-CoA (Zhang et al. 2005). Importantly, a human neurodegenerative disorder has been associated with a defect of the *PANK2* gene, encoding the mitochondrial isoenzyme in mammals (Hayflick 2014).

Similar to *CAB1*, genes encoding the remaining four enzymes of yeast CoA biosynthesis are also essential for cellular viability. 4’-Phosphopantothenate as the product of the PanK reaction is used for formation of 4’-phosphopantothenoylcysteine (PPC) which is subsequently decarboxylated to give 4’-phosphopantetheine (PP). While these biosynthetic steps are catalyzed by a bifunctional enzyme in *E. coli* (PPC synthetase PPCS and PPC decarboxylase PPCDC encoded by *coaBC*), individual enzymes exist in *S. cerevisiae* (encoded by *CAB2* and *CAB3*, respectively; Olzhausen et al. 2009) and higher eukaryotes (Daugherty et al. 2002). Exome sequencing in human patients gave evidence that PPCS variants may cause cardiomyopathy (Iuso et al. 2018). In addition to Cab3, yeast heterotrimeric PPCDC also contains subunits Hal3 (= Sis2) and Vhs3 which were initially described as negative regulators of phosphatase Ppz1 involved in halotolerance (Ruiz et al. 2009; Abrie et al. 2015). Thus, Hal3 and Vhs3 are “moonlighting” proteins at least one of which is essential for yeast viability. PP then reacts with the adenylyl moiety of ATP to give Dephospho-CoA (DPC) and finally DPC is phosphorylated, forming CoA. Yeast monofunctional enzymes PP adenylyltransferase (PPAT) and DPC kinase (DPCK) encoded by *CAB4* and *CAB5*,respectively (*coaD* and *coaE* in *E. coli*), are required to complete CoA biosynthesis while a bifunctional CoA synthase exists in mammals (Zhyvoloup et al. 2002). Mutations in the human *COASY* gene are associated with neurodegeneration (Dusi et al. 2014). Besides its enzymatic PPCDC activity, Cab3 physically interacts with Cab2, Hal3, Vhs3, Cab4 and Cab5 (but not with Cab1/PanK) and thus functions as a scaffold of the yeast CoA synthesizing protein complex (CoA-SPC) with a molecular weight of about 330 kDa (Olzhausen et al. 2013). Assembly of such a CoA biosynthetic complex has been recently also shown for mammalian cell lines (Bakovic et al. 2021). There is evidence that *Drosophila* and mice may use PP as an extracellular precursor which is used to complete biosynthesis of CoA in the absence of PANK, PPCS and PPCDC (Srinivasan et al. 2015; reviewed by Sibon and Strauss 2016). Fig. 1 summarizes the biosynthesis of CoA in *S. cerevisiae* and shows metabolic pathways requiring acetyl-CoA together with some applications in biotechnology.

**Fig. 1.**
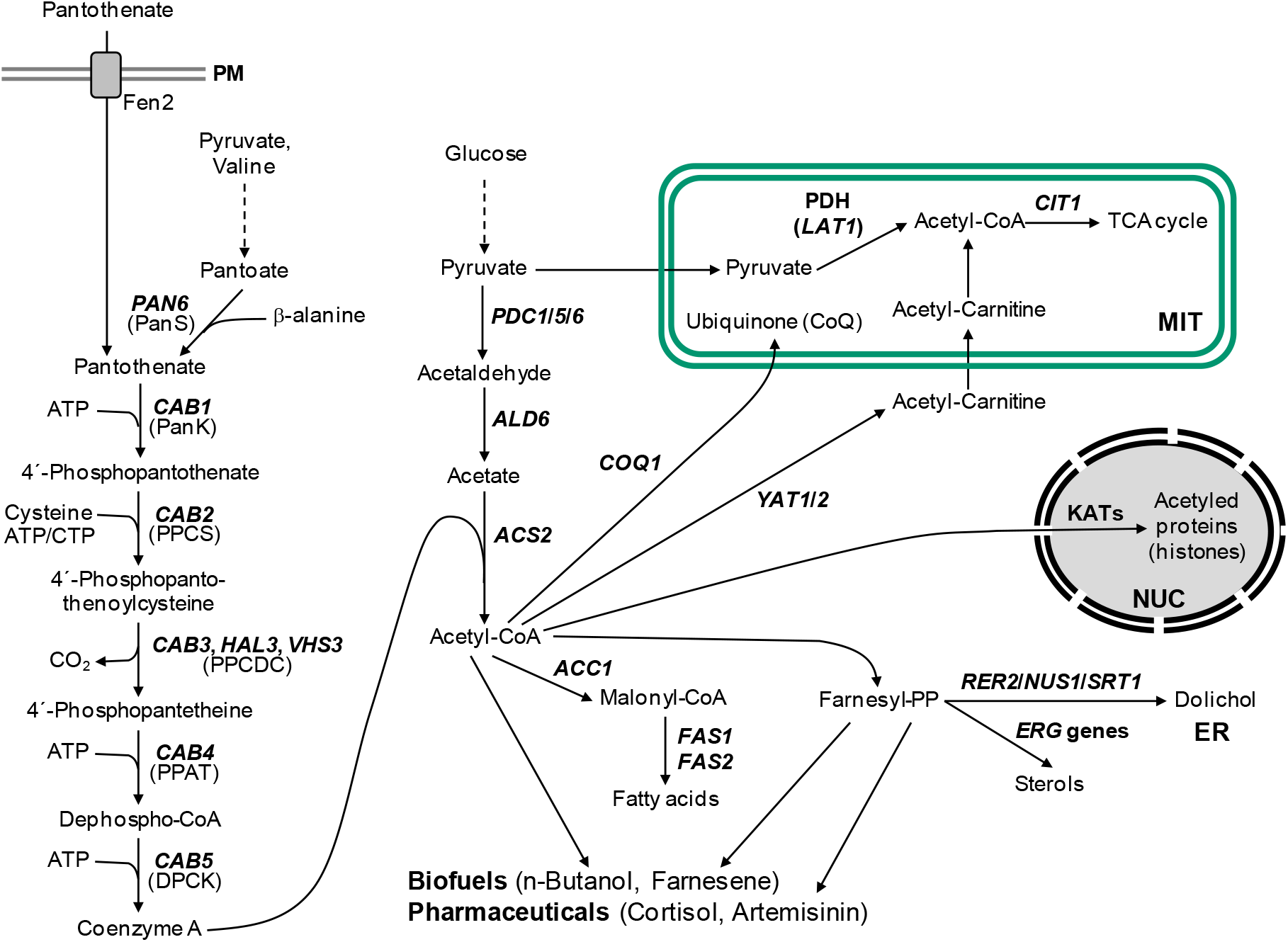
Biosynthesis of CoA in *S. cerevisiae*, metabolic pathways requiring acetyl-CoA and applications of acetyl-CoA in molecular biotechnology. Acetyl-CoA as a substrate of the glyoxylate cycle essential for utilization of C_2_-substrates (e. g. ethanol) is not shown. Abbreviations of cellular compartments: ER, endoplasmic reticulum; MIT, mitochondria; NUC, nucleus; PM, plasma membrane. Enzymes of CoA biosynthesis: PanS, pantothenate synthase; PanK, pantothenate kinase; PPCS, 4’-phosphopantothenoylcysteine synthetase; PPCDC, 4’-phosphopantothenoylcysteine decarboxylase; PPAT, 4’-phosphopantetheine adenylyltransferase; DPCK, Dephospho-CoA kinase.

Since all yeast *CAB* genes are expressed at a low level, gene overexpression may be considered as a means for engineering of the CoA biosynthetic pathway. However, it appears reasonable to assume that inhibition of PanK activity by the final product of the pathway prevents an increased metabolic flux towards CoA, as previously shown in *E. coli* (Song and Jackowski 1992). We thus devised a strategy to overcome the presumed PanK inhibition by CoA/acetyl-CoA and were indeed able to identify a hyperactive and constitutive enzyme variant (W331R). Stable overexpression of this variant in combination with additional genes of CoA biosynthesis and physiological modifications of growth media finally allowed us to construct yeast strains in which the intracellular concentration of CoA nucleotides was increased almost 40-fold above the wild-type level.

## Materials and methods

### Yeast strains and media

All strains of *S. cerevisiae* used in this work were derived from regulatory wild-type strains JS91.15-23 (Olzhausen et al. 2009) and BY4741 (Brachmann et al. 1998) by successive transformation with gene cassettes encoding *TPI1-CAB* fusions (see below). Complete genotypes of major strains are shown in Table 1 (an extended list of all strains and genotypes is available as Supplementary Table S1).

**Table 1:** Strains of *Saccharomyces cerevisiae* used in this work.

| Strain | Genotype |
| --- | --- |
| JS91.15-23 | *MATα ura3 leu2 trp1 his3* |
| BY4741 | *MATa ura3 leu2 his3 met15* |
| JS91.14-24 | *MATα ura3 his3 cab1<sup>ts</sup> (G351S)* |
| MGY22<br>(derived from JS91.15-23) | *MATα ura3 leu2 trp1 his3*<br>*HIP1::TPI1<sub>PRO</sub>-HA<sub>3</sub>-CAB2-CYC1<sub>TER</sub>*<br>*ENO1::TPI1<sub>PRO</sub>-HA<sub>3</sub>-CAB3-CYC1<sub>TER</sub>*<br>*MRH1::TPI1<sub>PRO</sub>-HA<sub>3</sub>-CAB4-CYC1<sub>TER</sub>*<br>*ADH3::TPI1<sub>PRO</sub>-HA<sub>3</sub>-CAB5-CYC1<sub>TER</sub>*<br>*ZPS1::TPI1<sub>PRO</sub>-HA<sub>3</sub>-CAB1<sub>W331R</sub>-CYC1<sub>TER</sub>*<br>*CUP9::TPI1<sub>PRO</sub>-HA<sub>3</sub>-HAL3<sub>aa 260-495</sub>-CYC1<sub>TER</sub>*<br>+ pSH62 (2μm *HIS3 GAL1-cre*) |
| MGY23<br>(derived from BY4741) | *MATa ura3 leu2 his3 met15*<br>*HIP1::TPI1<sub>PRO</sub>-HA<sub>3</sub>-CAB2-CYC1<sub>TER</sub>*<br>*ENO1::TPI1<sub>PRO</sub>-HA<sub>3</sub>-CAB3-CYC1<sub>TER</sub>*<br>*MRH1::TPI1<sub>PRO</sub>-HA<sub>3</sub>-CAB4-CYC1<sub>TER</sub>*<br>*ADH3::TPI1<sub>PRO</sub>-HA<sub>3</sub>-CAB5-CYC1<sub>TER</sub>*<br>*ZPS1::TPI1<sub>PRO</sub>-HA<sub>3</sub>-CAB1<sub>W331R</sub>-CYC1<sub>TER</sub>*<br>*CUP9::TPI1<sub>PRO</sub>-HA<sub>3</sub>-HAL3<sub>aa 260-495</sub>-CYC1<sub>TER</sub>*<br>+ pSH62 (2μm *HIS3 GAL1-cre*) |
| LRY2 | *MATα ura3 leu2 trp1 his3*<br>*HIP1::TPI1<sub>PRO</sub>-HA<sub>3</sub>-CAB2-CYC1<sub>TER</sub>*<br>*ENO1::TPI1<sub>PRO</sub>-HA<sub>3</sub>-CAB3-CYC1<sub>TER</sub>*<br>*MRH1::TPI1<sub>PRO</sub>-HA<sub>3</sub>-CAB4-CYC1<sub>TER</sub>*<br>*ADH3::TPI1<sub>PRO</sub>-HA<sub>3</sub>-CAB5-CYC1<sub>TER</sub>*<br>*ZPS1::TPI1<sub>PRO</sub>-HA<sub>3</sub>-CAB1<sub>W331R</sub>-CYC1<sub>TER</sub>*<br>*CUP9::TPI1<sub>PRO</sub>-HA<sub>3</sub>-HAL3<sub>aa 260-495</sub>-CYC1<sub>TER</sub>*<br>*GAT1::TPI1<sub>PRO</sub>-HA<sub>3</sub>-FEN2-CYC1<sub>TER</sub>*<br>*pcd1Δ::LEU2* + pSH62 (2μm *HIS3 GAL1-cre*) |

Synthetic complete (SC) media were prepared by using 0.17% Yeast Nitrogen Base as a source of vitamins (final concentration of pantothenate: 0.84 μM). Concentration of ammonium sulfate, amino acids and bases has been described (Olzhausen et al. 2009). Transformants were selected with SCD-Leu medium (2% glucose, without leucine). For improved supply of a pantothenate precursor, β-alanine was added to a final concentration of 200 mg/l (2.25 mM). Concentration of pantothenate was elevated 50-fold (20 mg/l; 42 μM) or 100-fold (40 mg/l; 84 μM) for certain cultivations.

### Plasmid constructions

Standard cloning vectors pRS415, pRS416 (Sikorski and Hieter 1989), p415-MET25, p416-MET25 and p426-MET25HA (Mumberg et al. 1994) were used for plasmid constructions employed in this work (Genetic markers are given in Supplementary Table S2).

Allelic variants of *CAB1* obtained by selection of conditional mutant JS91.14-24 (*cab1*^ts^ G351S) at 37°C were amplified by PCR using reading frame primers CAB1-Bam-Start + CAB1-Hind-Stop and chromosomal DNA prepared from suppressor strains obtained. Amplification products were cleaved with BamHI + HindIII and inserted into expression plasmid p416-MET25 to give various pLS12-S plasmids. As a wild-type reference, pTM8 was constructed accordingly.

Multi-copy expression plasmids pSBS5 (*CAB1*) and pJO73 (murine PanK3 gene) have been described (Olzhausen et al. 2009). Variants of *CAB1* for functional analysis were obtained by site-directed mutagenesis (see below) and inserted into p415-MET25 and p426-MET25HA.

The construction of plasmid pLEUTEX3 used for integration of CoA biosynthetic genes activated by the *TPI1* promoter is described in detail in the Supplementary Information. Coding regions of *CAB1, CAB1* W331R, *CAB2, CAB3, CAB4, CAB5, HAL3*_aa260-495_ and *FEN2* were amplified with gene-specific primers and individually inserted into the multi-cloning site of pLEUTEX3 to give *TPI1-CAB* fusions. For site-specific recombination with selected chromosomal loci, flanking sequences of about 400 bp were amplified and ligated to both sides of pLEUTEX plasmids. Recombination of gene expression cassettes with the desired genomic position was verified by analytical PCR, using chromosomal DNA of transformants and specific verification primers outside the flanking regions. After each transformation, the selection marker *loxP-LEU2*-loxP was removed by activation of the *GAL1*-dependent cre recombinase on plasmid pSH62 (cultivation in SCGal-His; Güldener et al. 1996).

### Site-directed mutagenesis

To introduce missense mutations into the *CAB1* coding region at selected sites, the QuikChange Site-Directed Mutagenesis Kit from Agilent was used. Plasmid pJO20 was derived from pUC19 by insertion of the *CAB1* coding region as a BamHI + HindIII fragment and subsequently used as a template for mutagenesis. Sequences of oligonucleotides used for mutagenesis are shown in Supplementary Table S3. After confirmation of the desired mutation by DNA sequencing, *CAB1* reading frame cassettes were transferred to various expression plasmids.

### Plasmid shuffling

Missense variants of *CAB1* and human *COASY* were functionally studied in *S. cerevisiae* using the plasmid shuffling strategy (Sikorski and Boeke 1991). Entire genes *CAB1, CAB4* and *CAB5*, respectively, were inserted into *ARS CEN URA3* vector pRS416 and the resulting rescue plasmids pJO57, pGE7 and pGE9 were individually transformed into wild-type strain JS91.15-23. Chromosomal copies of these *CAB* genes were subsequently deleted, using gene disruption cassettes released from plasmids pSBS7 (*cab1*Δ::*HIS3*), pSB2 (*cab4*Δ::*HIS3*) and pSB5 (*cab5*Δ::*HIS3*), respectively (Olzhausen et al. 2009). The resulting strain JS19.1 (*cab1*Δ::*HIS3* [*CAB1*]) was then transformed with *ARS CEN LEU2* plasmid pKH45 (*CAB1*, positive control) and related plasmids containing various missense mutations at selected positions. Similarly, the complementation of null mutations *cab4*Δ and *cab5*Δ by pGE8 (*CAB4*), pGE10 (*CAB5*) and pLS20 (h*COASY*) was tested by transformation of LSY20 (*cab4*Δ::*HIS3* [*CAB4*]) and LSY21 (*cab5*Δ::*HIS3* [*CAB5*]), respectively. Transformants were incubated on synthetic medium containing 5-fluoroorotic acid (FOA) to counter-select against *URA3* rescue plasmids.

### Enzyme assay

Assay of pantothenate kinase has been described (Olzhausen et al. 2009). In brief, the assay depends on ATP-dependent conversion of D-[1-^14^C] pantothenate (55 mCi/mMol; Biotrend, Cologne, Germany) to phosphopantothenate and its binding to DEAE cellulose ion exchange filter disks which were subsequently analyzed by scintillation counting (6 assays for each PanK variant). Since strain JS91.14-24 used as a recipient for transformation also contains a conditional PanK, assays were performed at 37°C.

To investigate the influence of CoA and its thioesters on the activity of PanK from yeast (Cab1) and mouse (PanK3 isoenzyme) in the absence of cellular metabolites, crude extracts from transformants of strain JS91.14-24 were partially purified by ion exchange chromatography. Prior to protein binding, DEAE sepharose CL-6B (Sigma-Aldrich) was equilibrated with binding buffer (100 mM Tris/HCl + 2.5 mM MgCl_2_, pH 7.4). Crude extracts (5 mg/ml protein) were incubated with DEAE sepharose at 4°C, washed 5 times with the 2.5-fold volume of binding buffer and finally treated with elution buffer containing 600 mM NaCl. Salt was removed by microdialysis against binding buffer. Inhibitory concentrations of CoA and its thioesters on PanK activity were calculated by using the GraphPad Prism software (San Diego, USA).

### Measurements of metabolites

Yeast strains were harvested at a density of 2 x 10^7^ cells/ml, collected by centrifugation and used for preparation of crude extracts by intensive agitation with glass beads for 6 min. Removal of proteins by precipitation with perchloric acid (final concentration: 400 mM), neutralization with K_2_CO_3_ and subsequent quantification of metabolites CoA and acetyl-CoA by enzyme-coupled reactions has been performed as described (Bergmeyer 1974). In brief, CoA and acetyl-CoA were assayed together by using the citrate synthase reaction which catalyzes the formation of citrate from acetyl-CoA and oxaloacetate. Oxaloacetate is provided by the NAD-dependent malate dehydrogenase reaction which also serves as the photometric indicator reaction (increase of absorbance by NADH at 340 nm). All CoA is converted to acetyl-CoA by phosphotransacetylase with acetylphosphate as a substrate. To distinguish between CoA and acetyl-CoA, N-ethylmaleimide was used to block the free SH group of CoA. Validity of the assay was verified by analysis of CoA + acetyl-CoA samples of known concentration and by adding defined concentrations of coenzymes to yeast crude extracts. Amounts of CoA + acetyl-CoA were calculated with respect to protein concentrations determined prior to protein removal from crude extracts. Routinely, each strain was cultivated three times and cell extracts derived were assayed in triplicate. Details on the assay conditions are provided as Supplementary Method.

### Miscellaneous procedures

For amplification of *CAB* genes by PCR, proofreading-competent *Pwo* DNA polymerase was used. Westernblot analysis of epitope-tagged PanK variants was performed with peroxidase-conjugated anti-HA antibody (Roche Diagnostics) and luminol-containing detection system. *CAB1* inserts of plasmids obtained by selection for second-site suppressor mutations of the *cab1*G351S allele or site-directed mutagenesis were completely sequenced to confirm the presence of the desired mutations and the absence of unwanted alterations (performed by LGC Genomics, Berlin, Germany).

## Results

### Intragenic suppressor mutations of a temperature-sensitive *cab1* mutant allele

In a previous work we could show that the mutant allele *cab1* G351S encodes a temperature-sensitive variant of *S. cerevisiae* pantothenate kinase (Olzhausen et al. 2009). Interestingly, overexpression of a *MET25-cab1* G351S promoter fusion using a multi-copy plasmid led to slow but substantial growth of transformants (not shown). Because of this residual activity we hypothesized that selection for growth of strain JS91.14-24 (chromosomal *cab1*^ts^ G351S mutation) at the nonpermissive temperature might allow identification of intragenic second-site mutations which reconstitute PanK enzyme activity. 30 revertants which could grow at 37°C on rich medium (YEPD) as well as on synthetic medium (SCD) similar to a wild-type strain were selected for further analysis. Chromosomal DNA was prepared from these strains and used for amplification of the *CAB1* coding region. After fusing these *CAB1* alleles with the *MET25* promoter of a single-copy vector, resulting plasmids were introduced into the *cab1*^ts^ strain. 27 plasmids could complement this mutation, indicating that they contain an intragenic suppressor mutation which again enables biosynthesis of a functional PanK. Comparative sequence analysis of the cloned *CAB1* variants showed that they represent four distinct intragenic second-site mutations (A22G, F103V, D114E and W331L). As shown in Fig. 2, transformants of the *cab1*^ts^ G351S mutant strain containing additional *CAB1* variants A22G G351S, F103V G351S, D114E G351S and W331L G351S exhibit a substantial increase of PanK activity (with W331L G351S being most effective) and were able to grow at 37°C similar to the wild-type. We conclude that variants A22G, F103V, D114 and W331L are critical for PanK activity of the yeast enzyme.

**Fig. 2.**
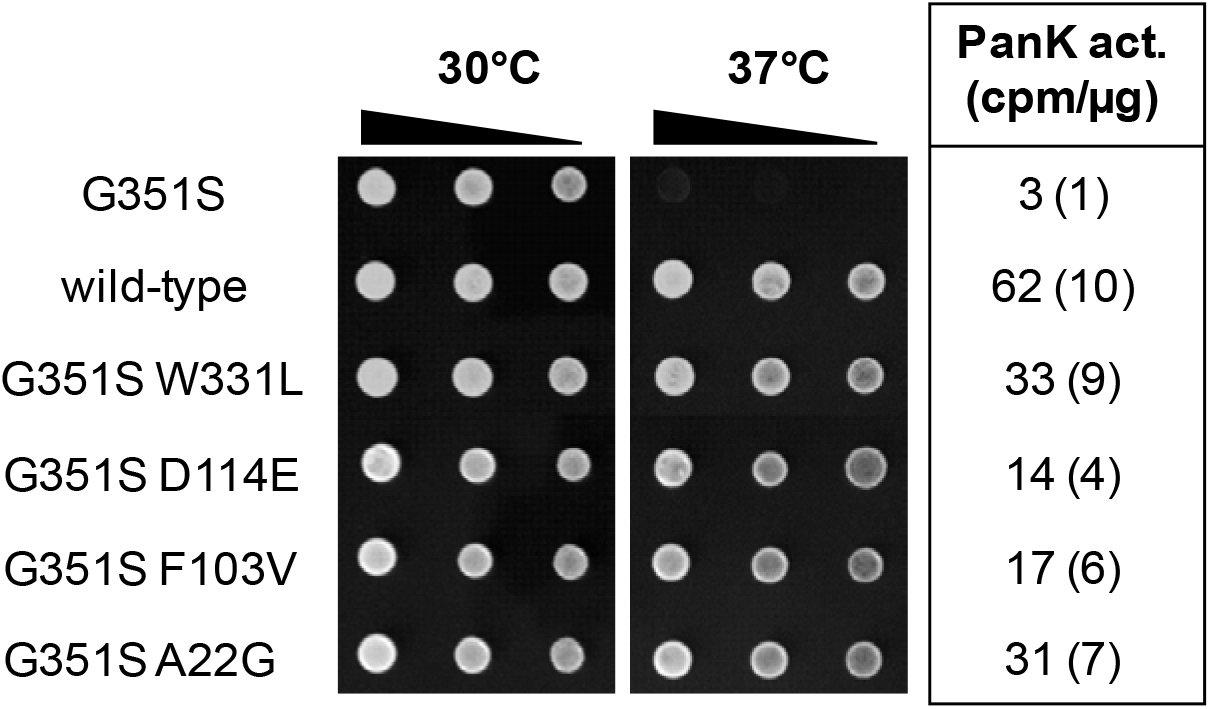
Phenotypic analysis of intragenic *cab1* G351S revertants. Single-copy plasmid p416-MET25 (empty vector, negative control) and expression plasmids containing *CAB1* wild-type (pTM8) and allele variants G351S W331L (pLS12-S8), G351S D114E (pLS12-S9), G351S F103V (pLS12-S10) and G351S A22G (pLS12-Y6) were transformed into strain JS91.14-24 (*ura3 cab1*^ts^ G351S). Serial dilutions of transformants were spotted on selective synthetic medium (SCD) and incubated at 30°C and 37°C, respectively. Pantothenate kinase (PanK) activity in protein extracts prepared from transformants was assayed at 37°C. For each assay, 75 μg of total protein was used. PanK activities are given in cpm 1-^14^C-phosphopantothenate formed per μg protein. Standard deviations are shown in parentheses.

### Inhibition of *S. cerevisiae* PanK by acetyl-CoA

It has been reported that the enzyme activity of eukaryotic pantothenate kinases from fungal and mammalian organisms is mainly regulated by acetyl-CoA and less effective by CoA (Calder et al. 1999; Zhang et al. 2005). We thus investigated whether CoA and its thioesters influence the activity of the *CAB1* encoded PanK from *S. cerevisiae*. Strain JS91.14-24 was transformed with a multi-copy expression plasmid containing a *MET25-CAB1* fusion and subsequently used to prepare partially purified PanK from a protein extract. PanK assays were performed at 37°C in the presence of varying concentrations of CoA and its thioesters acetyl-CoA, malonyl-CoA and palmitoyl-CoA. As is apparent from Fig. 3b, acetyl-CoA clearly inhibits PanK activity (IC_50_ = 36 μM; inhibitory concentration leading to 50% reduction of PanK activity). In contrast, some inhibition was also observed with CoA, malonyl-CoA and palmitoyl-CoA but significantly higher concentration were needed (IC_50_ = 283 μM, 812 μM and 891 μM, respectively; Fig. 3a, c, d).

**Fig. 3.**
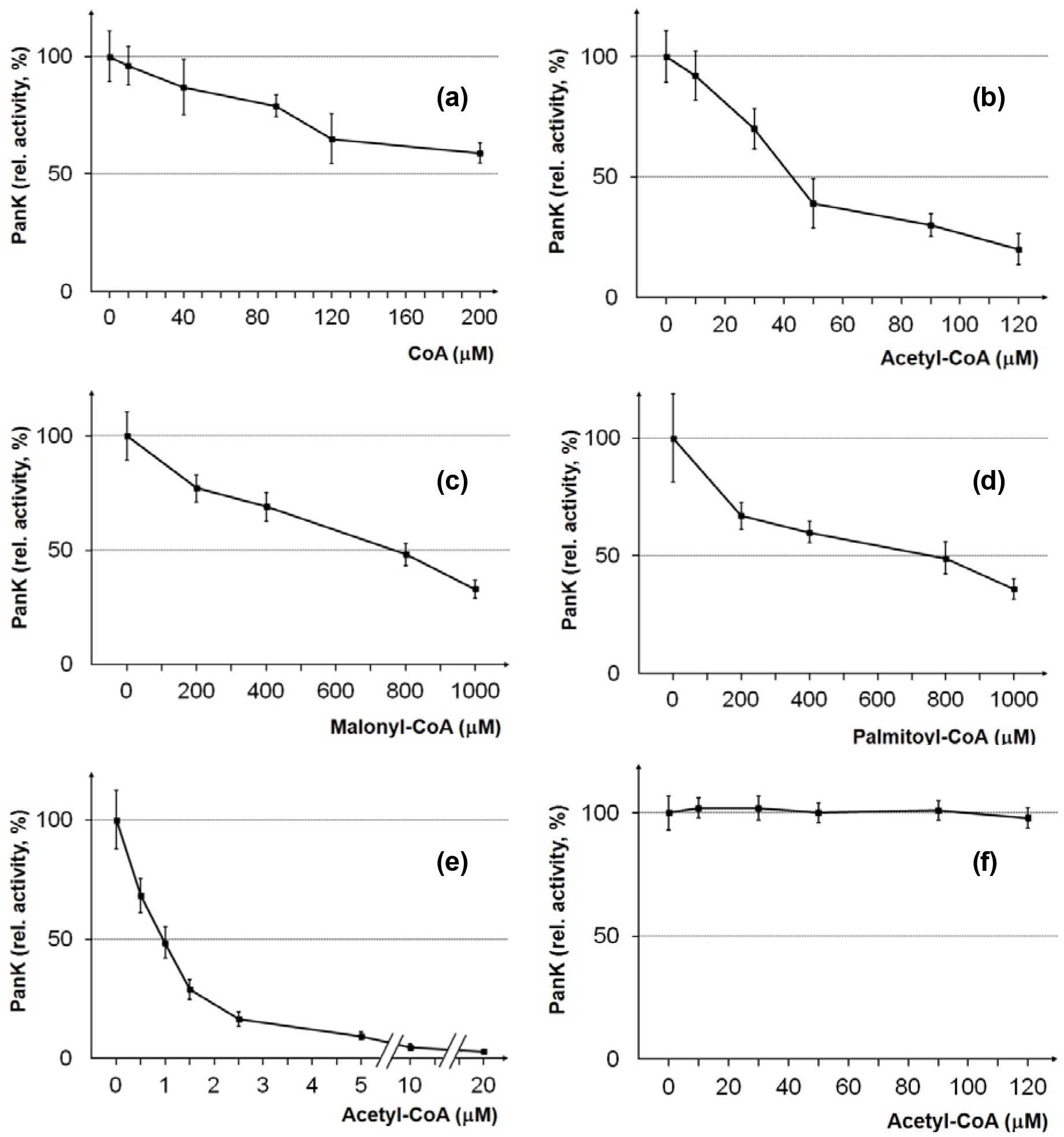
Influence of CoA and its acyl derivatives on pantothenate kinases. Multi-copy expression plasmids pSBS5 (*MET25-CAB1*; wild-type; a-d), pJO73 (*MET25-PANK3*; murine wild-type; e) and pEB27 (*MET25-CAB1* W331R; hyperactive yeast allele; f) were transformed into strain JS91.14-24 (*cab1*^ts^). To remove contaminating soluble yeast metabolites, crude extracts of transformants were partially purified by PanK binding to DEAE sepharose, elution and subsequent dialysis. Varying concentrations of CoA (a), acetyl-CoA (b, e, f), malonyl-CoA (c) and palmitoyl-CoA (d) as indicated were added to PanK assay mixtures. Relative PanK activities (%) refer to enzyme activities in the absence of CoA inhibitors (100%). Standard deviations for at least four independent assays per inhibitor concentration are shown.

Previously, inhibitory concentrations for acetyl-CoA of murine isoenzymes PanK1β (IC_50_ = 10 μM) and PanK3 (IC_50_ = 1 μM; Zhang et al. 2005) have been reported. For a direct comparison with the yeast enzyme, we thus studied acetyl-CoA-dependent activity of PanK3 from *Mus musculus* (IC_50_ = 1 μM; Fig. 3e) and could demonstrate that the single PanK from *S. cerevisiae* is clearly less sensitive against inhibition by acetyl-CoA than mammalian PanK3.

### Site-directed mutagenesis of the PanK acetyl-CoA binding region

Inhibition of PanK activity by acetyl-CoA must be considered as a key mechanism regulating the CoA biosynthetic pathway, thus preventing overproduction of CoA. We thus reasoned that overexpression of *CAB* genes may not be sufficient to increase the cellular CoA level. However, this problem may be overcome by construction of a PanK variant which is no longer affected by acetyl-CoA.

To obtain such a PanK variant, we took advantage of the crystal structure obtained for the human PanK3 isoenzyme in the presence of acetyl-CoA (Hong et al. 2007). Structural modelling indicated that binding sites of pantothenate as a substrate and acetyl-CoA as a competitive feedback inhibitor partially overlap (cf. Supplementary Fig. S1). While several amino acid side chains are presumably required for interaction with substrate and inhibitor, some residues may be specifically involved in binding of acetyl-CoA. Thus, it should be possible to generate PanK variants of the wild-type which are not affected for binding of pantothenate but do no longer interact with acetyl-CoA, uncoupling binding of substrate and inhibitor. Following alignment of human PanK3 and yeast Cab1 sequences we selected residues for site-directed mutagenesis presumably important for binding of pantothenate, ATP and/or acetyl-CoA (N155, S158, R173, Y326, F330, W331, A233 and I234). It should be emphasized that W331 forming hydrophobic interactions with the methyl group of acetyl-CoA has been independently identified by selecting suppressors of the *cab1*^ts^ mutant allele G351S (see above). *CAB1* variant W331L was constructed because it corresponds to the second-site suppressor of the G351S allele but now in the sequence context of an intact enzyme. We also constructed a W331R variant which should abolish binding of acetyl-CoA even more effectively. Arginine instead of tryptophan was selected because the PanK from *Staphylococcus aureus* (*coaA* gene product) which is moderately related to the yeast enzyme (18% identity, 29% similarity; cf. Supplementary Fig. S2) but insensitive to CoA and acetyl-CoA (Leonardi et al. 2005a) contains this amino acid (R244) at the position corresponding to yeast W331 (W340 in human PanK3).

*CAB1* reading frame cassettes containing the desired mutations were inserted into a multi-copy expression plasmid, activated by the *MET25* promoter. Protein extracts from transformants of strain JS91.14-24 (*ura3 cab1*^ts^) were used to assay PanK activities. Using the plasmid shuffling strategy (Sikorski and Boeke 1991), *CAB1* variants were also tested for functional complementation of a *cab1* null mutation. We thus transformed a wild-type strain with a single-copy *URA3* plasmid containing the authentic *CAB1* gene, deleted the chromosomal copy of *CAB1* and subsequently introduced single-copy *LEU2* plasmids encoding *CAB1* variants. Use of a synthetic medium containing FOA (5-fluoroorotic acid) allows for counter-selection of plasmids with *URA3* as a selection marker so that growth entirely depends on the functionality of *CAB1* variants present on *LEU2* plasmids.

*CAB1* variants S158V, R173A and I234E failed to complement a *cab1* null mutation (no growth on FOA-containing synthetic medium; Fig. 4a) and encoded PanK enzymes had completely or almost completely lost their activity (Fig. 4b). All gene variants tested could be stably expressed in strain JS91.14-24 (Fig. 4c). Variants N155V and A233E were able to complement the *cab1* null allele and showed residual PanK activity (31% and 22% of the wild-type level, respectively). Loss of aromatic residues affecting interaction with the acetyl moiety of acetyl-CoA (double variant Y326A F330A and W331L) led to slight or substantial increase of PanK activity. Importantly, Cab1 W331R turned out as a hyperactive PanK variant the activity of which was more than 4-fold increased compared with the wild-type enzyme. We finally investigated whether the activity of this PanK variant responds to acetyl-CoA. In contrast to the wild-type enzyme, Cab1 W331R was completely insensitive to increased concentrations of acetyl-CoA (compare Fig. 3b and Fig. 3f). We conclude that the hyperactive and completely deregulated PanK variant W331R may be a suitable tool for engineering the CoA biosynthetic pathway.

**Fig. 4.**
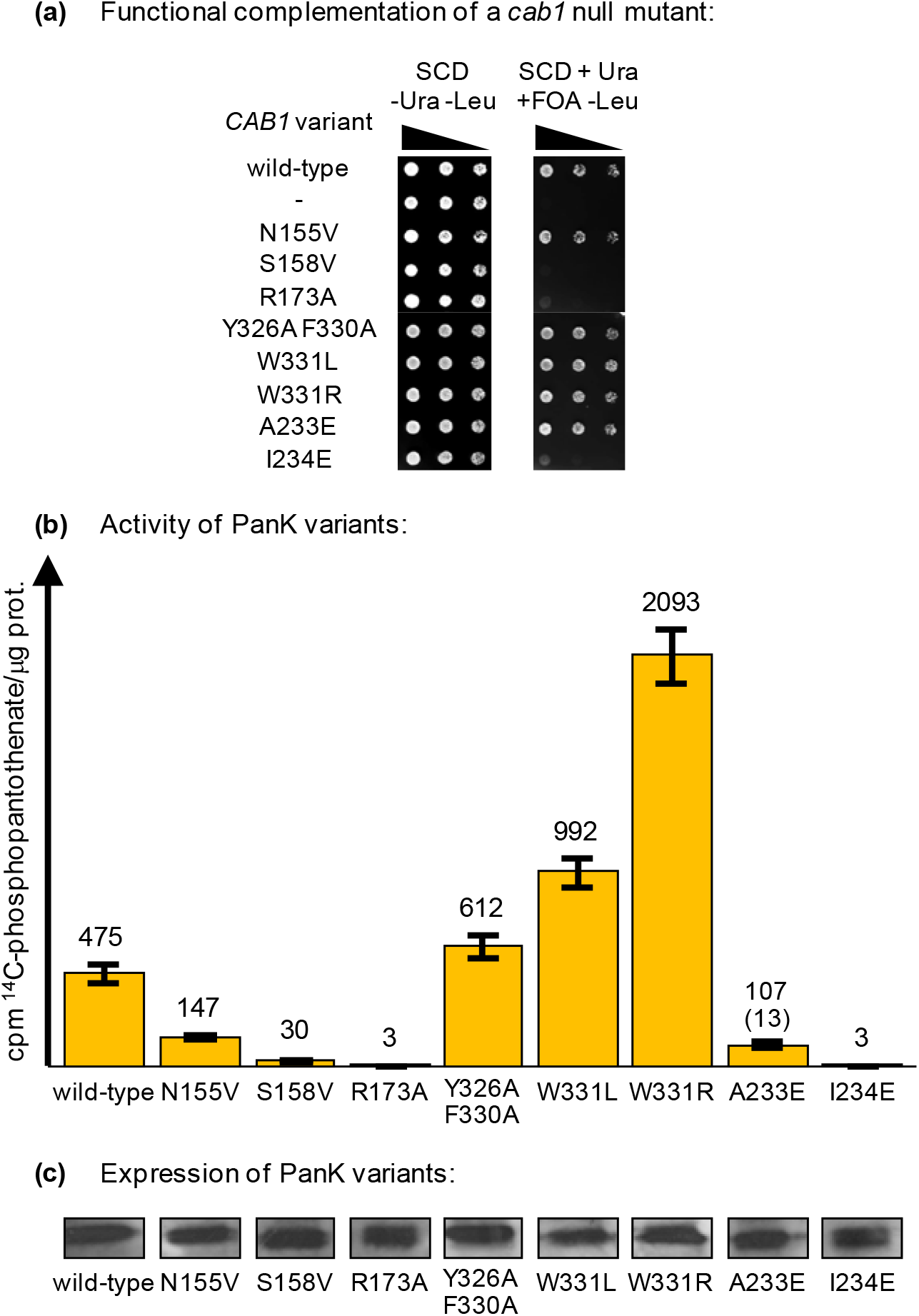
Functional analysis of *CAB1* variants. (a) Strain JS19.1 with a chromosomal *cab1*Δ null allele and *ARS CEN URA3 CAB1* rescue plasmid (pJO57) was transformed with *ARS CEN LEU2* plasmids containing the *CAB1* variants indicated. All transformants were able to grow on double-selective synthetic medium (SCD-Ura-Leu) while only functional variants of *CAB1* allowed growth on FOA-containing medium after loss of the rescue plasmid. FOA: 5-Fluoroorotic acid. (b) Activity of PanK variants constructed by site-directed mutagenesis. The coding region of wild-type *CAB1* was mutagenized at the positions indicated and gene variants were inserted into a multi-copy expression vector containing the *MET25* promoter together with the HA epitope. The resulting plasmids pSBS5 (*CAB1* wild-type), pEB5 (*CAB1* Y326A F330A), pEB6 (*cab1* I234E), pEB8 (*CAB1* W331L), pEB22 (*CAB1* N155V), pEB23 (*cab1* S158V), pEB25 (*cab1* R173A), pEB26 (*CAB1* A233E) and pEB27 (*CAB1* W331R) were transformed into strain JS91.14-24 (*cab1*^ts^). Transformants were cultivated at 30°C until the mid-log growth phase. Pantothenate kinase (PanK) activity in protein extracts prepared from transformants was assayed at 37°C. For each assay, 75 μg of total protein was used. PanK activities are given in cpm 1-^14^C-phosphopantothenate formed per μg protein. Standard deviations are indicated by error bars. (c) Stable expression of PanK variants was investigated by Western blot analysis of protein extracts prepared from transformants using anti-HA antibodies.

### Functional analysis of human *COASY* in *S. cerevisiae*

To facilitate pathway engineering of yeast CoA biosynthesis we supposed that the mammalian gene *COASY* encoding the bifunctional CoA Synthase could be possibly used instead of individual genes *CAB4* and *CAB5*. Assaying *COASY* for functional complementation of null mutations *cab4* and *cab5* should provide evidence whether such a strategy is helpful. For this analysis, we again used the strategy of plasmid shuffling (as described above for functional studies of *CAB1* variants). Wild-type strains with single-copy *URA3* rescue plasmids containing *CAB4* or *CAB5* were used to introduce null mutant alleles *cab4*Δ::*HIS3* and *cab5*Δ::*HIS3*, respectively. To assay for functional complementation, a cDNA encoding the human *COASY* gene (564 aa) driven by the yeast *MET25* promoter was inserted into a single-copy *LEU2* plasmid. Corresponding *LEU2* plasmids containing authentic yeast genes *CAB4* and *CAB5*, respectively, were constructed and used as a positive control. As shown in Fig. 4a, *hCOASY* could partially complement a *cab4*Δ mutation (although clearly less effective than *CAB4*) while only residual growth was observed with *hCOASY* in the *cab5*Δ mutant strain. Human CoA synthase could be efficiently synthesized in *S. cerevisiae* (Fig. 5b), indicating that this result is not a consequence of failure to express the heterologous gene. Since full-length hCoasy is associated with mitochondria (Zhyvoloup *et al*., 2003), we repeated this experiment with a variant lacking aa 1-29 being responsible for this targeting in mammalian cell lines. However, the truncated *hCOASY* variant encoding aa 30-564 was completely unable to complement null mutations *cab4*Δ and *cab5*Δ, respectively (not shown). We thus conclude that the bifunctional COASY gene cannot be used to simplify engineering of the CoA biosynthetic pathway in yeast. A similar strategy can be considered with bacterial *coaBC*, encoding a bifunctional enzyme corresponding to Cab2 + Cab3. Although *coaBC* could complement null mutations *cab2* and *cab3*, transformants showed slower growth, compared with authentic genes *CAB2* and *CAB3*, respectively (not shown).

**Fig. 5.**
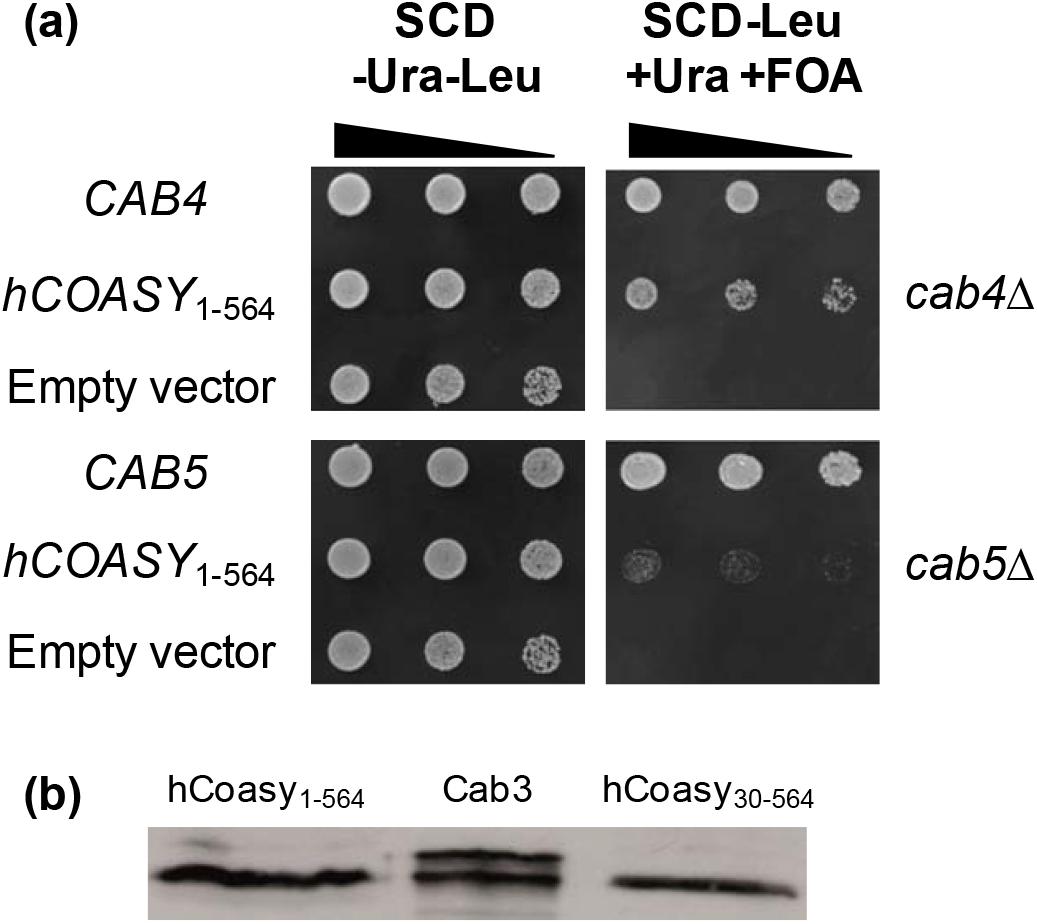
Functional analysis of human CoA Synthase gene (*COASY*) in *S. cerevisiae*. (a) For plasmid shuffling, strains LSY20 (*cab4*Δ + rescue plasmid pGE7 [*ARS CEN URA3 CAB4*]; upper part) and LSY21 (*cab5*Δ + rescue plasmid pGE9 [*ARS CEN URA3 CAB5*]; lower part) were transformed with the *ARS CEN LEU2* plasmid pLS20 containing the human *COASY* gene activated by the *MET25* promoter. Plasmids pGE8 (*CAB4*) and pGE10 (*CAB5*) served as positive controls, empty vector YCp111 as a negative control. FOA: 5-Fluoroorotic acid. (b) Stable biosynthesis of full-length hCoasy and truncated variant in transformants of *S. cerevisiae*. Expression plasmids pLS14 and pLS15 encoding HA-tagged length variants hCoasy_1-564_ and hCoasy_30-564_, respectively, were transformed into strain JS91.15-23. Protein extracts were analyzed by immuno-blotting using anti-HA antibodies.

### Construction of strains stably overexpressing CoA biosynthetic genes

To elevate the cellular level of CoA and/or acetyl-CoA, we wished to overproduce all enzymes being important for this pathway. This should be achieved by stable integration of additional gene copies, activated by a strong control region. Our previous characterization of *CAB* genes showed that these genes are expressed at a moderate or low level and that expression is not substantially influenced by the carbon source or availability of amino acids (Olzhausen et al. 2009). Consequently, use of an effective control region to elevate *CAB* gene transcription in combination with the allele *CAB1* W331R may allow increasing the biosynthesis of CoA and/or acetyl-CoA which now should no longer inhibit the initial PanK reaction of the pathway. We thus selected the strong *TPI1* promoter of the glycolytic enzyme triosephosphate isomerase for high-level expression of genes *CAB1* (wild-type and W331R variant), *CAB2*, *CAB3*, *HAL3*, *CAB4*, *CAB5* and *FEN2*. Although PPCDC is a heterotrimeric enzyme in yeast, co-overexpression of *CAB3* and *HAL3* should be sufficient for a substantially elevated enzyme activity (Ruiz et al. 2009). Since Hal3 is a moonlighting enzyme with an additional metabolic function, only aa 260-495 of Hal3 which are relevant for its PPCDC function (Hal3^PD^; Abrie et al. 2012) were overproduced. To study the possible influence of increased supply of pantothenate, we decided to also overexpress *FEN2*, encoding the pantothenate transporter of the plasma membrane (Stolz and Sauer 1999).

For stable chromosomal integration of up to seven additional genes, we constructed plasmid pLEUTEX3, containing *LEU2* as a selection marker flanked by two loxP sites. Thus, *LEU2* can be removed later by induction of the cre recombinase (marker rescue). The plasmid also contains *TPI1* promoter and *CYC1* terminator, separated by the HA epitope and a versatile cloning site for insertion of genes of CoA biosynthesis. Finally, flanking sequences from a suitable chromosomal site must be added to enable integration of the expression cassette at a position of choice by homologous recombination. The functional elements of this cassette are summarized in Fig. 6. Details of the construction procedure are provided as Supplementary description. Because of the mobile *LEU2* selection marker, repeated integration events of *TPI1-CAB* gene fusions are possible.

**Fig. 6.**
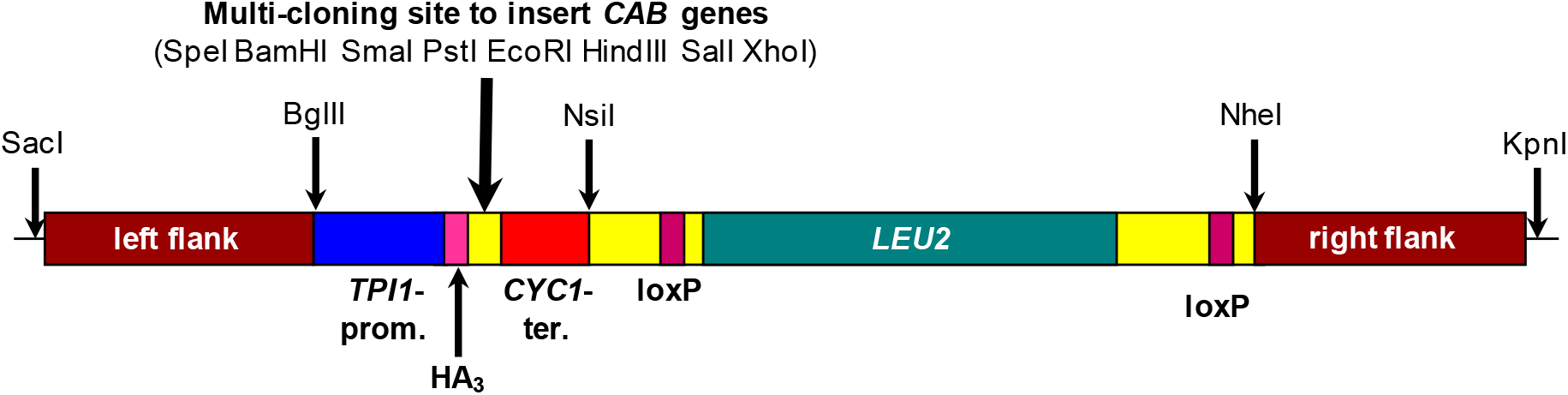
Functional elements of pLEUTEX expression cassettes suitable for repeated integration of CoA biosynthetic genes. Selection marker *LEU2* flanked by loxP sites can be removed after a successful transformation by induction of cre recombinase which targets loxP sites. Several unique restriction sites following the *TPI1*-HA promoter/epitope segment allow insertion of *CAB* reading frame cassettes. For integration of the expression/selection cassette by homologous recombination, flanking sequences from a genomic site of choice must be inserted by using restriction sites *Sac*I + *Bgl*II (left flank) and *Nhe*I + *Kpn*I (right flank), respectively. Prior to yeast transformation, expression cassettes together with flanking regions can be released by cleavage with *Sac*I + *Kpn*I.

Using basic plasmid pLEUTEX3, a number of expression cassettes with individual coding regions of CoA biosynthetic genes activated by *TPI1* were constructed. For subsequent site-specific integration, flanking sequences from genes with an extended upstream region were selected. Thus, positions of integration should not interfere with expression of nearby genes, preventing a possibly reduced fitness of transformants with additional *CAB* genes. Two upstream fragments of about 400 bp from selected genomic sites were amplified by PCR and ligated to the expression cassettes on both sides (selected positions:,, upstream“ regions of genes *ADH3, CUP9, ENO1, GAT1, HIP1, MRH1* und *ZPS1*). Release of the complete linear cassette including its flanking regions by cleavage with restriction enzymes and subsequent transformation of a *leu2* mutant should allow its stable integration by homologous recombination at the desired gene locus (Rothstein 1991). Plasmids derived from pLEUTEX3 which were subsequently used to integrate additional copies of CoA biosynthetic genes are listed in Table 2.

**Table 2:** Plasmids for targeting of expression cassettes at selected genomic sites.

| Plasmid | Genetic markers |
| --- | --- |
| pLEUTEX-CAB1 | *ZPS1*-5'-flank::*TPI1*<sub>PRO</sub>-HA<sub>3</sub>-*CAB1*-*CYC1*<sub>TER</sub>::*ZPS1*-3'-flank |
| pLEUTEX-CAB1<br>W331R | *ZPS1*-5'-flank::*TPI1*<sub>PRO</sub>-HA<sub>3</sub>-*CAB1*<sub>W331R</sub>-*CYC1*<sub>TER</sub>::*ZPS1*-3'-flank |
| pLEUTEX-CAB2 | *HIP1*-5'-flank::*TPI1*<sub>PRO</sub>-HA<sub>3</sub>-*CAB2*-*CYC1*<sub>TER</sub>::*HIP1*-3'-flank |
| pLEUTEX-CAB3 | *ENO1*-5'-flank::*TPI1*<sub>PRO</sub>-HA<sub>3</sub>-*CAB3*-*CYC1*<sub>TER</sub>::*ENO1*-3'-flank |
| pLEUTEX-CAB4 | *MRH1*-5'-flank::*TPI1*<sub>PRO</sub>-HA<sub>3</sub>-*CAB4*-*CYC1*<sub>TER</sub>::*MRH1*-3'-flank |
| pLEUTEX-CAB5 | *ADH3*-5'-flank::*TPI1*<sub>PRO</sub>-HA<sub>3</sub>-*CAB5*-*CYC1*<sub>TER</sub>::*ADH3*-3'-flank |
| pLEUTEX-HAL3 | *CUP9*-5'-flank::*TPI1*<sub>PRO</sub>-HA<sub>3</sub>-*HAL3*<sub>aa260-495</sub>-*CYC1*<sub>TER</sub>::*CUP9*-3'-flank |
| pLEUTEX-FEN2 | *GAT1*-5'-flank::*TPI1*<sub>PRO</sub>-HA<sub>3</sub>-*FEN2*-*CYC1*<sub>TER</sub>::*GAT1*-3'-flank |

In addition, all plasmids carry *LEU2* as a selection marker together with loxP sequences (loxP::*LEU2*::loxP) between *CYC1*<sub>TER</sub> and the 3'-flank.

### Increased production of acetyl-CoA by overexpression of *CAB* genes

After having completed the construction of 8 pLEUTEX plasmids (7 different genes and *CAB1* wild-type and W331R variant, respectively), linear expression constructs were integrated one after another into *leu2* strains JS91.15-23 and BY4741, representing different strain backgrounds. Our previous work on CoA biosynthesis was performed with strain JS91.15-23 (Olzhausen et al. 2009) while BY4741 has been routinely used for functional gene studies (Brachmann et al. 1998). To investigate whether such differences possibly influence our intended engineering of CoA biosynthesis, both strains were treated in parallel. The order of integration events (*CAB2* – *CAB3* – *CAB4* – *CAB5* – *CAB1* [wild-type or W331R variant] – *HAL3* – *FEN2*) is apparent from the complete list of constructed strains (cf. Table 1 of Supplementary Information). Successful integration of each expression construct at the correct genomic positions was verified by analytic PCR with gene-specific primers (not shown). Transformants also contained episomal plasmid pSH62, encoding a *GAL1-cre* fusion which allowed galactose-inducible synthesis of cre recombinase for removal of loxP-flanked selection marker *LEU2* after each step (Güldener et al. 1996).

Strains MGY22 (derived from JS91.15-23) and MGY23 (derived from BY4741) were obtained by multiple rounds of gene integration and contain additional genes *CAB1* W331R, *CAB2*, *CAB3*, *HAL3*, *CAB4* and *CAB5*. Both strains (together with their isogenic reference strains without additional *CAB* genes) were cultivated in YPD rich medium until mid-log phase and subsequently used for preparation of protein-free extracts. To quantify CoA biosynthesis in these metabolite extracts, an enzymatic assay simultaneously recording CoA + acetyl-CoA was used (Bergmeyer 1974). To distinguish between both nucleotides, NEM (N-ethylmaleimide) was added to certain assays, blocking the free SH group of CoA. As a result, only acetyl-CoA is able to react in the assay system.

Importantly, strong overexpression of 6 CoA biosynthetic genes (*CAB1* W331R *CAB2 CAB3 HAL3 CAB4 CAB5*) resulted in a 15-fold increase of CoA nucleotides in the JS strain background when transformants were grown in YPD rich medium (0.9 μMol/g protein in the reference strain vs. 13.5 μMol/g protein in strain MGY22 with additional *CAB* genes; Fig. 7). Almost identical results were obtained with transformants in the BY background (17-fold increase compared with the reference strain). The level of CoA nucleotides was slightly reduced after NEM had been added to extracts from the reference strain (regular *CAB* gene dosage), indicating that most CoA exists as acetyl-CoA which is insensitive against NEM. This finding appears plausible, considering the inhibition of PanK by acetyl-CoA and much weaker by CoA (see above, Fig. 3a, b). In the overproducing strain MGY22, acetyl-CoA constitutes 78% of total CoA. Interestingly, cultivation of strain MGY22 in synthetic SCD medium led to a substantially lower level of CoA nucleotides as found after growth in YPD (2.6-fold increase vs. 15-fold increase; Fig. 7). In contrast, CoA biosynthesis in the SCD-grown reference strain is similar to the same strain grown in YPD. This finding provides evidence for a limitation of precursor molecules when the overproducing strain is grown in standard SCD, preventing a level of CoA biosynthesis as observed in rich medium.

**Fig. 7.**
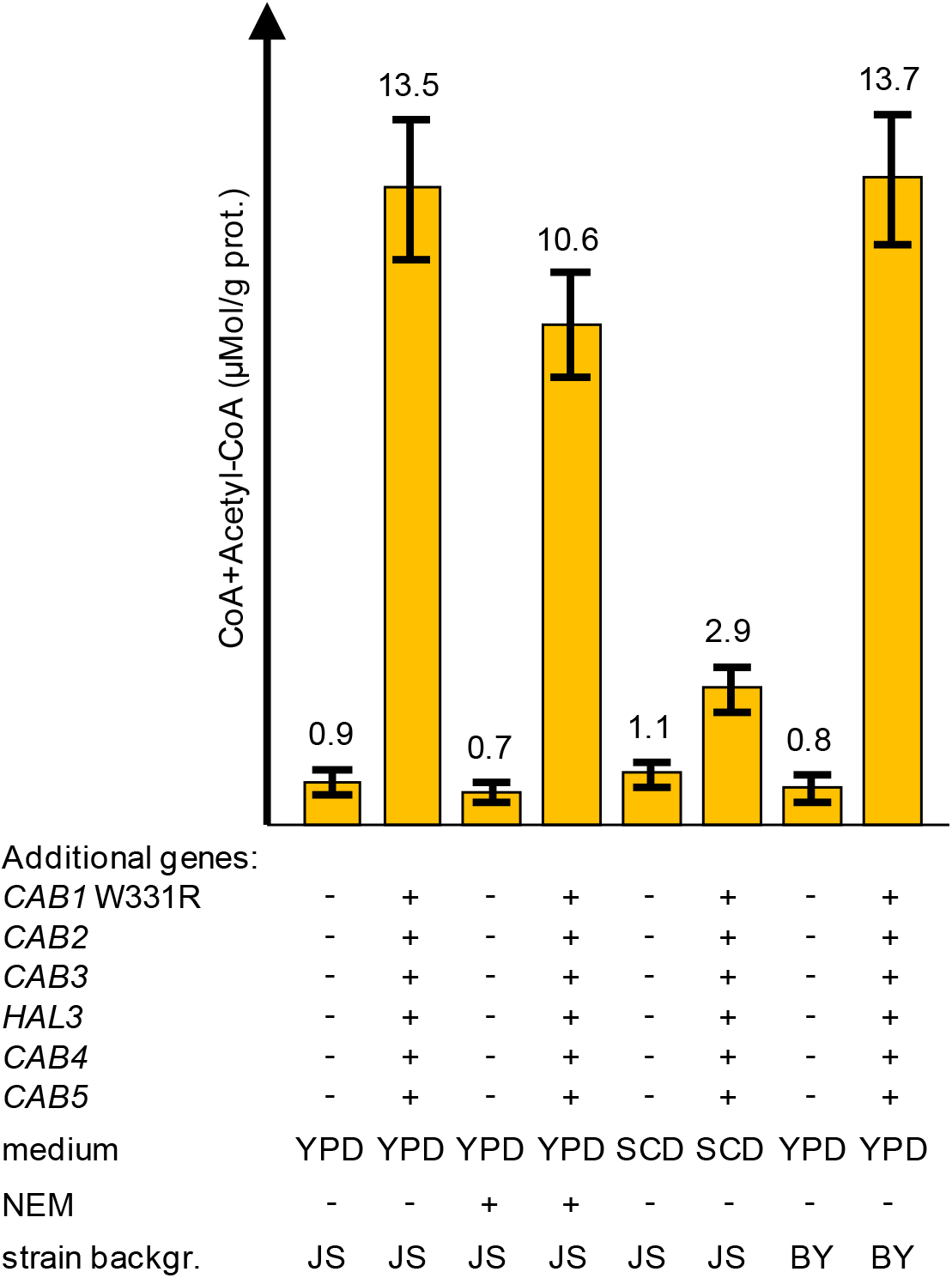
Influence of *CAB* gene dosage variation on biosynthesis of CoA nucleotides. CoA and acetyl-CoA were simultaneously measured by enzymatic analysis in protein-free cell extracts of reference strains JS91.15-23 / BY4741 and transformants MGY22 / MGY23. Additional *TPI1*-dependent genes introduced into strains are indicated by +. Transformants were cultivated in rich medium (YPD) or synthetic complete medium (SCD), both containing 2% glucose as a carbon source. NEM (N-ethylmaleimide) was added to certain reactions to distinguish between CoA and acetyl-CoA. For comparison, transformants of two different strain backgrounds (JS and BY) were investigated. Standard deviations are indicated by error bars.

As an estimation of *CAB* gene overexpression by *TPI1*-dependent transcription, we comparatively assayed PanK activity in strains JS91.15-23 (regular gene dosage) and MGY22 (additional *TPI1-CAB* genes). It turned out that the activity of pantothenate kinase in protein extracts of MGY22 was about 13-fold increased relative to the reference (JS91.15-23: 72 cpm/μg protein, MGY22: 950 cpm/μg protein).

### Influence of genetic and physiological variations on production of acetyl-CoA

After having shown that overexpression of CoA biosynthetic genes led to a strong increase of acetyl-CoA in protein-free cell extracts, we wished to study the relative contribution of the genes involved in more detail. Since engineering of the CoA biosynthetic pathway led to similar results in both strain backgrounds tested, subsequent studies focused on strain JS91.15-23 and its derivatives. As described above, variant *CAB1* W331R present in strain MGY22 encodes a PanK which is completely insensitive against inhibition by acetyl-CoA. We thus investigated whether overexpression of wild-type *CAB1* is able to similarly increase CoA biosynthesis as observed with the deregulated variant. As is shown in Fig. 8, *TPI1*-dependent expression of wild-type *CAB1* in combination with 5 additional genes of CoA biosynthesis (strain MGY24) was also able to increase the production of CoA + acetyl-CoA (4.1-fold, compared with the reference strain). However, this increase remained substantially below the level obtained in the presence of *CAB1* W331R (15-fold increase), proving that release of PanK from inhibition by acetyl-CoA is an essential prerequisite for efficient engineering of CoA biosynthesis. We next studied whether gene variant *CAB1* W331R may be solely responsible for elevation of CoA nucleotides. Indeed, strain TUY1 containing only the *TPI1-CAB1* W331R expression cassette and regular dosage of the remaining CoA biosynthetic genes showed a clearly increased level of CoA + acetyl-CoA (4.8-fold increase) which nevertheless was lower than the yield observed with strain MGY22. We also investigated the influence of PPCDC subunits on CoA biosynthesis, comparing two strains with and without a *TPI1-HAL3* expression cassette. In strain MGY18 devoid of this cassette, CoA + acetyl-CoA were 13-fold elevated with respect to the reference (Fig. 8), arguing for only a minor contribution of additional Hal3 to the CoA biosynthetic pathway.

**Fig. 8.**
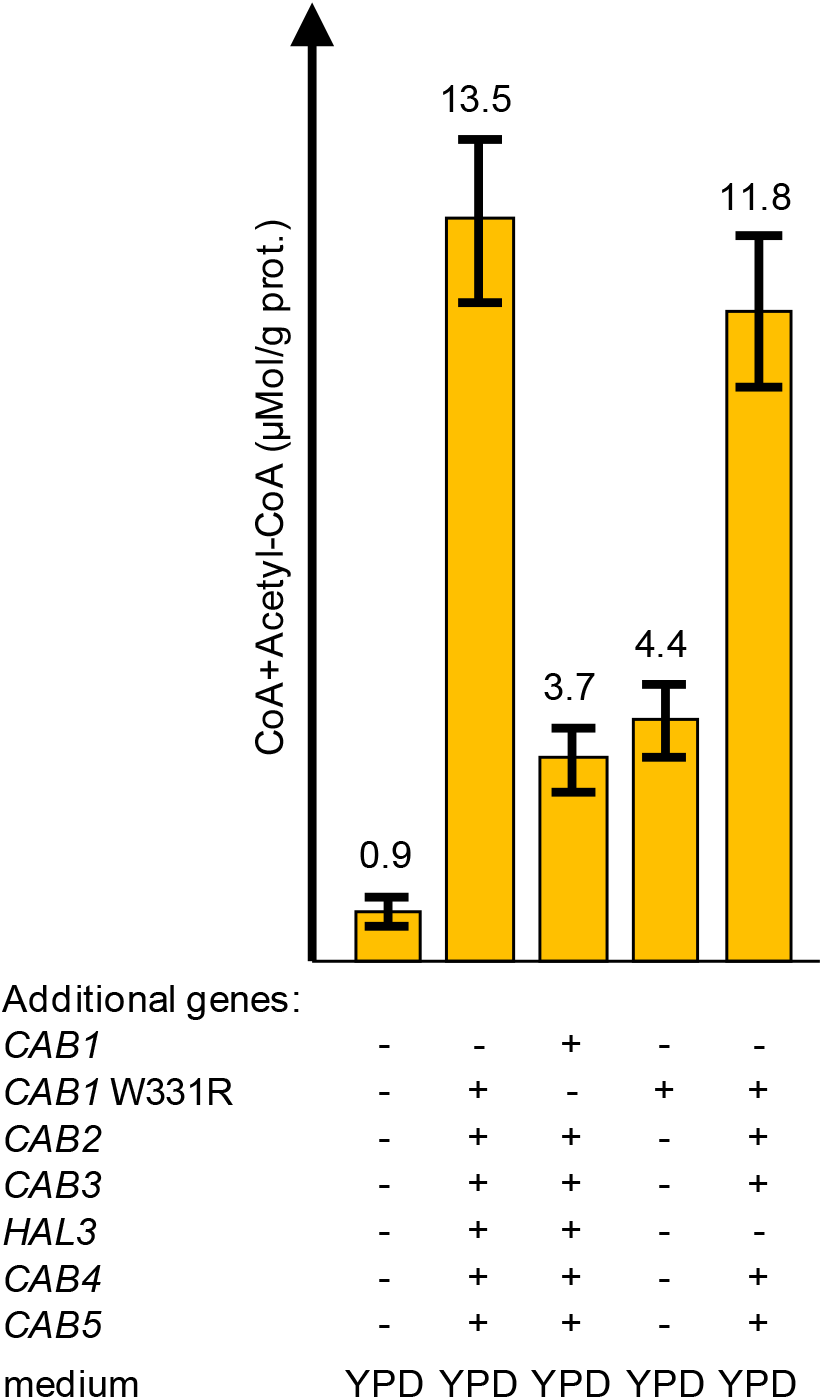
Various combinations of additional *CAB* genes and influence on biosynthesis of CoA nucleotides. CoA and acetyl-CoA were simultaneously measured by enzymatic analysis in protein-free cell extracts of reference strain JS91.15-23 and transformants MGY22 / MGY24 / TUY1 / MGY18. Additional *TPI1*-dependent genes introduced into strains are indicated by +. All transformants were cultivated in rich medium (YPD). Standard deviations are indicated by error bars.

Our finding that production of CoA nucleotides in MGY22 after cultivation in standard synthetic medium (SCD) was substantially below the level observed with YPD rich medium strongly indicates limited supply with a critical nutrient. We thus supplemented standard SCD with precursors β-alanine (200 mg/l) or pantothenate (20 mg/l; 50-fold increase relative to the contribution of yeast nitrogen base) which can be taken up by plasma membrane transporters Gap1 (general amino acid permease) and Fen2 (Sauer and Stolz 1999), respectively. While imported pantothenate can be directly used by PanK, β-alanine must react with pantoate (catalyzed by pantothenate synthase, *PAN6*) to finally give pantothenate. The availability of β-alanine may be critical for pantothenate biosynthesis because yeast, in contrast to *E. coli*, does not contain an aspartate decarboxylase gene and generates β-alanine inefficiently from polyamines (White et al. 2001).

Interestingly, even in the unmodified reference strain, supplementation of SCD medium with β-alanine or pantothenate could elevate the amount of CoA + acetyl-CoA (2.3 and 4.8-fold; Fig. 9). Growth of strain MGY22 in SCD supplemented with β-alanine stimulated CoA biosynthesis about 4-fold while additional pantothenate increased the amount of CoA nucleotides almost 10-fold (28.4 μMol/g protein; Fig. 9), exceeding even the productivity observed after cultivation in YPD rich medium (13.5 μMol/g protein; Fig. 7). Although the concentration of pantothenate in rich medium is unknown, it may be still limiting. Indeed, supplementation of YPD with additional 20 mg/l pantothenate resulted in a further increase of CoA + acetyl-CoA (30.7 μMol/g protein; Fig. 9). Together, these results confirm that the availability of pantothenate is a critical factor for the metabolic flux towards CoA.

**Fig. 9.**
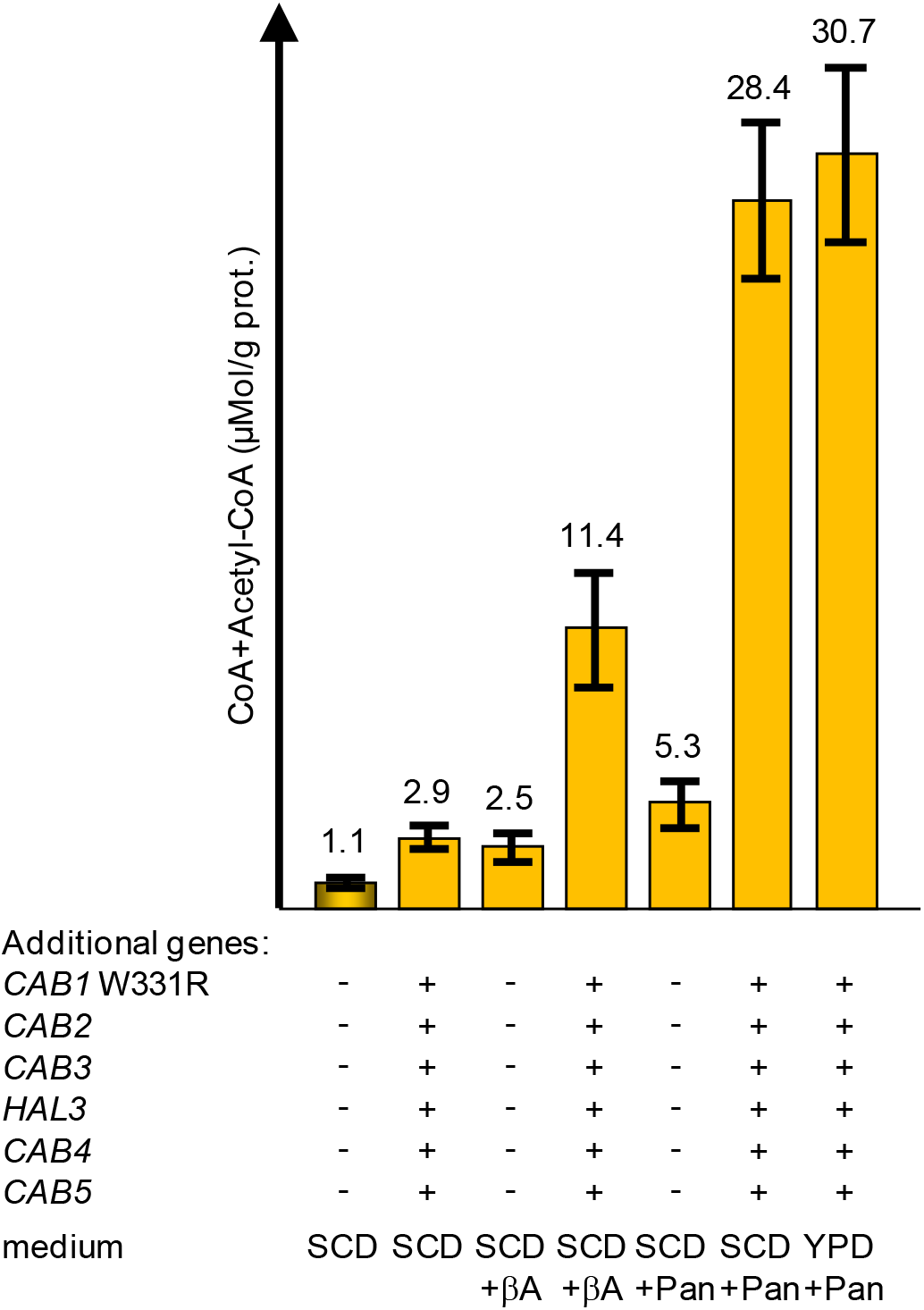
Influence of media supplementation on biosynthesis of CoA nucleotides. CoA and acetyl-CoA were simultaneously measured by enzymatic analysis in protein-free cell extracts of reference strain JS91.15-23 and transformant MGY22. Additional *TPI1*-dependent genes introduced into MGY22 are indicated by +. Transformants were cultivated in synthetic complete medium (SCD; 0.4 mg/l Ca-pantothenate; 0.84 μM) or rich medium (YPD), both containing 2% glucose as a carbon source. Media were supplemented to final concentrations of 200 mg/l β-alanine (βA; 2.25 mM) and 20 mg/l Ca-pantothenate (Pan, 50-fold compared with standard-SCD, 42 μM), respectively. Standard deviations are indicated by error bars.

Assuming that CoA biosynthesis can be further improved by an additional copy of the pantothenate permease gene, we finally decided to introduce a *TPI1-FEN2* expression cassette into MGY22. The resulting strain TUY4 was then cultivated in media enriched with pantothenate (50- and 100-fold, respectively). As is shown in Fig. 10, increase of the *FEN2* gene dosage could indeed further elevate the concentration of CoA nucleotides when pantothenate was present in non-limiting amounts. This increase was more apparent when transformants MGY22 and TUY4 were cultivated with 100-fold pantothenate (40 mg/l; + 27%) compared with 50-fold pantothenate (20 mg/l; + 16%). Supplementation with even higher concentrations of pantothenate could not substantially elevate the cellular yield of CoA nucleotides (not shown).

**Fig. 10.**
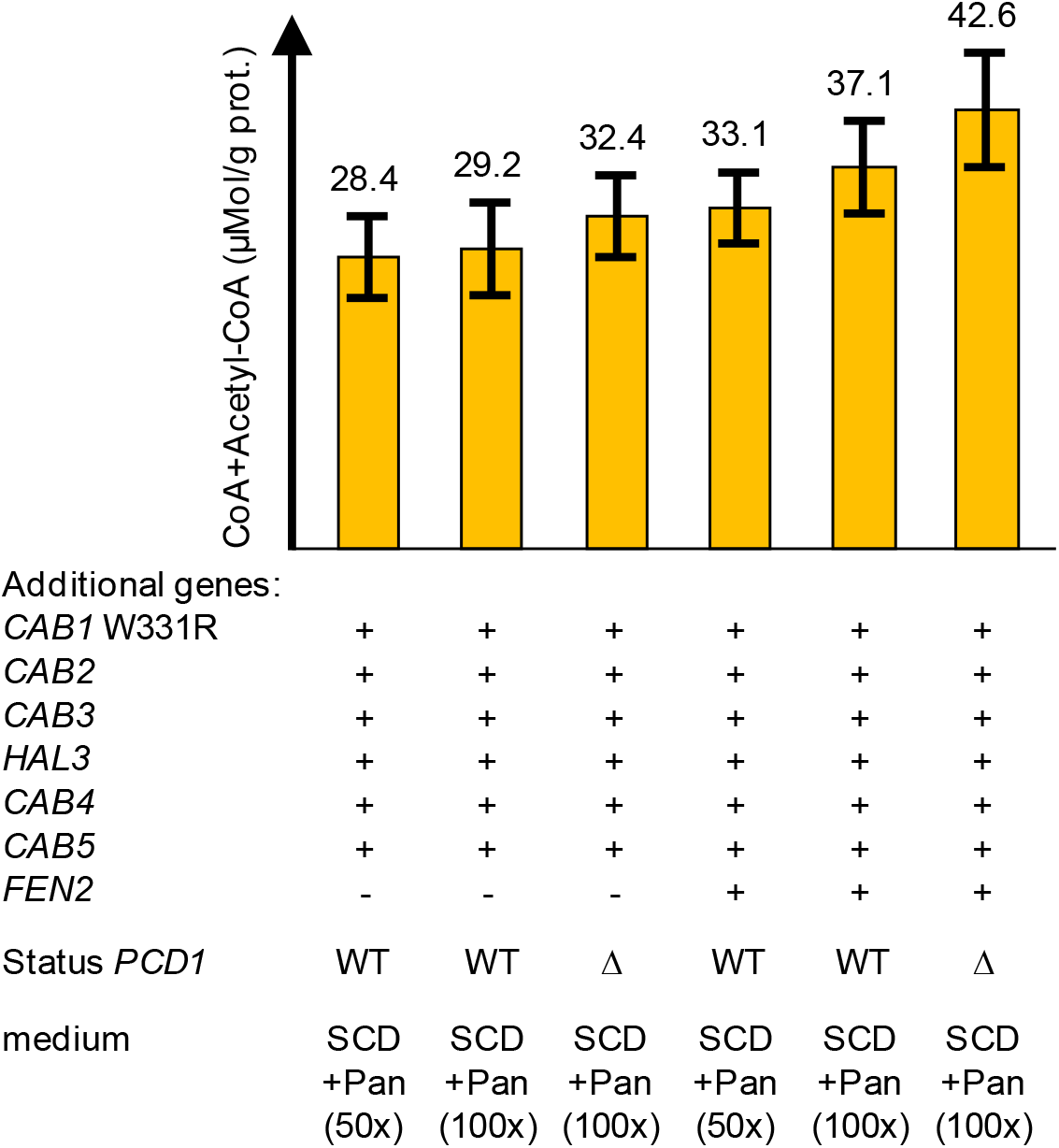
Influence of *FEN2* overexpression and deletion of *PCD1* on biosynthesis of CoA nucleotides. CoA and acetyl-CoA were simultaneously measured by enzymatic analysis in protein-free cell extracts of transformants MGY22, TUY4 (+ *FEN2*), LRY1 (Δ*pcd1*) and LRY2 (+ *FEN2* Δ*pcd1*). Additional *TPI1*-dependent genes are indicated by +. Transformants were cultivated in synthetic complete medium (SCD), containing 2% glucose as a carbon source. Media were supplemented to final concentrations of 20 mg/l or 40 mg/l Ca-pantothenate (Pan, 50- or 100-fold compared with standard-SCD, 42 μM or 84 μM). Deletion of the genomic *PCD1* gene encoding a CoA phosphatase is shown by Δ. Standard deviations are indicated by error bars.

The cellular level of CoA may not only be affected by its biosynthesis but also by degradation of the nucleotide structure. The nudix hydrolase encoded by *PCD1* encodes a pyrophosphatase which is effective against oxidized derivatives of CoA (CoA disulfide) but the purified enzyme also showed a substantial specificity for CoA and acetyl-CoA (Cartwright et al. 2000). Thus, we finally introduced a *pcd1*Δ null mutant allele into strains MGY22 and TUY4 (to give LRY1 and LRY2, respectively) and comparably investigated whether the concentration of CoA nucleotides changed. Indeed, with both strains we observed a further increase of CoA + acetyl-CoA (by 11-15%; Fig. 10), indicating that some hydrolysis even of intact CoA occurs when *PCD1* is functional.

## Discussion

Engineering of a pathway not only requires overexpression of the biosynthetic genes involved but has also taken into account that activity of enzymes may be affected by feedback inhibition by the end-product of the pathway. Similar to what has been shown for *E. coli* (Rock et al. 2003) this work demonstrates that biochemical regulation of PanK is a key mechanism controlling the metabolic flux towards CoA and its major thioester derivative acetyl-CoA. Taking advantage of the crystal structure of *E. coli* CoaA in the presence of the feedback inhibitor CoA, Rock et al. (2003) constructed and characterized a CoaA variant (R106A) which exhibited 54% activity of the wild-type enzyme but was completely insensitive against inhibition by CoA. As a result, the intracellular level of CoA increased about fourfold. For a mammalian cell line transfected with a *mPANK1*β expression construct, a CoA level 13 times higher than with control cells was reported (Rock et al. 2000).

In this work we show that the *S. cerevisiae* PanK (Cab1) is inhibited by acetyl-CoA with an IC_50_ concentration similar to what has been reported for the enzyme from *A. nidulans* (36 μM for ScPanK and 32 μM for AnPanK; Calder et al. 1999). We used a dual strategy to identify variants of Cab1 which are no longer inhibited by acetyl-CoA: (I) Genetic selection for intragenic revertants of the conditional mutant *cab1* G351S with restored growth at 37°C led to identification of the double mutation G351S W331L. PanK activity of the encoded enzyme reached 53% of the wild-type level, explaining why the growth defect of the single mutation could be suppressed. We hypothesized that a mutation improving activity of a defective enzyme may also positively affect the activity of the wild-type enzyme. (II) W331 is a strictly conserved residue of eukaryotic pantothenate kinases involved in binding of acetyl-CoA (Hong et al. 2007). Release from competitive inhibition of PanK may be achieved by replacing amino acids which specifically influence binding of acetyl-CoA but leave substrate binding sites unaffected. We thus replaced amino acids involved in inhibitor binding by residues presumed to be non-functional and assayed activity and regulation of the resulting yeast PanK variants. Some of these variants turned out to be partially (N155V, A233E) or totally defective (S158V, R173A, I234E), indicating that binding of substrates is severely impaired. However, Cab1 variants with the double mutation Y326A F330A or the single mutation W331L exhibited elevated PanK activities, confirming that it is indeed possible to uncouple binding of inhibitor and substrates. Sequence alignment of Cab1 and the unregulated CoaA from *S. aureus* revealed the existence of an arginine residue at the position corresponding to W331 which allowed us to construct W331R as a yeast PanK which was even more active than the variant W331L identified by our suppressor selection. Importantly, variant W331R increased the activity of PanK by a factor of four and rendered the enzyme completely insensitive against inhibition by acetyl-CoA. The observed hyperactivity indicates that PanK variant W331R not only prevents binding of the inhibitor acetyl-CoA but may also facilitate substrate access. It should be mentioned that Hong et al. (2007) obtained variants of human PanK3 some of which also showed elevated enzyme activity (e. g. 1.85-fold increase of PanK3 R78C). However, variants characterized by Hong et al. (2007) affect enzyme positions different from our mutations introduced into Cab1. Additional suppressor mutations of the *cab1* G351S mutant allele (A22G, F103V and D114E) were not further characterized in this work because the corresponding positions are not conserved among eukaryotic PanK enzymes. In addition, the crystal structure of human PanK3 failed to provide evidence for a function in inhibitor binding (Hong et al. 2007).

Although the CoA biosynthetic pathway is universal, structural organization of the enzymes involved may differ. This is apparent for the bifunctional mammalian enzyme COASY, corresponding to individual gene products Cab4 and Cab5 in *S. cerevisiae*. For our intended engineering of CoA biosynthesis we thus hypothesized that the *COASY* gene may be a convenient substitute for *CAB4* and *CAB5* to increase the metabolic flux through the final biosynthetic steps. However, it turned out that *COASY* could only partially complement a *cab4* null mutation and was unable to replace Cab5. Consequently, for the intended engineering of CoA biosynthesis only authentic genes from *S. cerevisiae* were used. Although the acetyl-CoA sensitive PanK encoded by *CAB1* is clearly of major regulatory importance we nevertheless decided to introduce additional copies of at least 6 genes (*CAB1* W331R, *CAB2, CAB3, HAL3, CAB4* and *CAB5*) overexpression of which should guarantee substantially elevated activities of 5 enzymes, thus preventing possible bottle-necks in the course of CoA biosynthesis. This was achieved by fusing *CAB* coding regions with the strong control region of the glycolytic gene *TPI1* and subsequent stable integration of fusion genes at selected genomic positions. Comparative PanK assays confirmed that the activity of this enzyme was indeed significantly elevated (13-fold). To avoid limitations with the number of fusion genes to be transformed into *S. cerevisiae* strains, we used a marker rescue strategy, employing *LEU2* flanked by loxP sites as a selection marker which allows its removal after each successful transformation step by induction of cre recombinase and repeated use of *LEU2* for further gene integrations.

Successive transformation of two *S. cerevisiae* strains representing different genetic backgrounds with 6 expression cassettes, cultivation of the resulting transformants in rich medium and quantification of CoA + acetyl-CoA in protein-free extracts revealed that these CoA nucleotides were indeed strongly overproduced in both strains at a similar level (MGY22: 15-fold; MGY23: 17-fold). As supposed, *CAB1* variant W331R encoding a hyperactive PanK insensitive against inhibition by acetyl-CoA substantially contributed to this increase. This was shown by comparative analysis of a strain identical to MGY22 but overexpressing the *CAB1* wild-type gene (MGY24; 4-fold increase). Although the *CAB1* W331R allele is of major importance for an elevated level of CoA nucleotides, the remaining *CAB* genes must be also overexpressed to achieve maximal overproduction (as shown by strain TUY1 with merely an extra copy of *CAB1* W331R; 4.8-fold overproduction).

Comparative cultivation of the CoA overproducer MGY22 in different media revealed that growth in synthetic medium SCD was unable to ensure an increased level of CoA nucleotides as observed after growth in rich medium YPD (2.6-fold increase vs. 15-fold). This deficiency could be overcome by supplementation of SCD with β-alanine or, even more effective, with pantothenate as the immediate precursor of the pathway, leading to a level of CoA nucleotides even higher as observed after growth in YPD. Indeed, pantothenate turned out as limiting in rich medium as well. We could also show that the presence of an additional *HAL3* gene supports biosynthesis of CoA + acetyl-CoA, presumably to guarantee full PPCDC activity together with *CAB3*. Previously, the importance of pantothenate as a “metabolic switch” influencing pathways which require CoA has been described (Sandoval et al. 2014). In the yeast *Schizosaccharomyces pombe*, overexpression of the *liz1^+^* gene homologous to the high-affinity H^+^-pantothenate symporter *FEN2* of *S. cerevisiae* could elevate the transport capacity of pantothenate 430-fold above the wild-type level (Stolz et al. 2004). In addition to 6 extra copies of CoA biosynthetic genes, we thus also introduced a *TPI1*-*FEN2* fusion into strain MGY22. Cultivating the resulting strain TUY4 in media heavily supplemented with pantothenate (50- and 100-fold), CoA nucleotides indeed showed a further increase (by 16% and 27%, respectively).

The intracellular concentration of CoA nucleotides may not only be influenced by their biosynthesis but also by degradation. Presumably, hydrolysis by a phosphatase (Pcd1 in yeast) is the initial step for complete degradation of CoA, finally releasing ADP, pantothenate and cysteamine (Naquet et al. 2020). Although Pcd1 has been described as a peroxisomal enzyme important for hydrolysis of oxidized CoA variants (such as CoA disulfide), degradation of intact CoA metabolites can also occur (at least in vitro with purified enzyme; Cartwright et al. 2000). Introduction of a *pcd1*Δ null allele into strains MGY22 and TUY4 again elevated the level of CoA nucleotides, confirming degradation as an important regulatory step which contributes to the CoA steady-state level. When strain LRY2 (overexpression of 7 additional genes with a *pcd1*Δ null mutation) was cultivated in pantothenate-supplemented SCD we were able to detect the highest level of CoA nucleotides found in the course of this work (42.6 μMol/g protein) which means an almost 39-fold increase, compared with the unmodified reference strain grown in standard SCD.

Because of this substantial overproduction described, our strains may be helpful tools to improve the metabolic flux towards biosynthetic pathways depending on acetyl-CoA. Phenotypic characterization of CoA-overproducing strains revealed that they are fully viable and are able to utilize carbon sources glucose, ethanol or acetate with the same efficiency as the wild-type. No morphological differences concerning cell size and budding pattern could be observed by microscopic inspection.

Previously, other strategies for improvement of biosynthetic pathways dependent on acetyl-CoA were used. Hong et al. (2019) overexpressed *CAB1* (regulatory wild-type) + *CAB3* + *ATF1* (encoding an alcohol acetyl transferase) to increase the yield of ethyl acetate and other esters in *S. cerevisiae*. For improved biosynthesis of the biofuel n-butanol by *S. cerevisiae*, Schadeweg and Boles (2016) decided to overproduce pantothenate kinase from *E. coli* (which is inhibited by CoA, but not by acetyl-CoA; *MET25-coaA* fusion) or the pantothenate permease (*MET25-FEN2* fusion). Indeed, both strategies were able to improve n-butanol production. In other work, introduction of bacterial genes encoding subunits of a cytosolic pyruvate dehydrogenase (without mitochondrial localization sequences) into a strain devoid of alcohol dehydrogenase genes (*adh*Δ) allowed a direct conversion of glycolytic pyruvate to acetyl-CoA, leading to an increased yield of n-butanol (Lian et al. 2014). The level of n-butanol could be also elevated after heterologous expression of the ATP-dependent citrate lyase gene *ACL1* from the oleaginous yeast *Yarrowia lipolytica*, enabling to cleave citrate into oxaloacetate and acetyl-CoA. Chen et al. (2013) were able to improve conversion of acetaldehyde into acetyl-CoA by co-expression of *ALD6* (encoding NADP-dependent aldehyde dehydrogenase) and a bacterial acetyl-CoA synthetase gene (*acs*SE^L641P^ of *Salmonella enterica*; resistant against inhibition by acetylation). While Hong et al. (2019) and Schadeweg and Boles (2016) partially intervened into the CoA biosynthetic pathway, these latter strategies increased the level of acetyl-CoA by shifting the ratio among CoA nucleotides without affecting total CoA concentration. Since the genetic manipulations described in this work affect all steps of CoA biosynthesis, the resulting substantial elevation especially of the acetyl-CoA concentration could be advantageous for existing and future strategies of metabolic engineering in *S. cerevisiae*.

## Acknowledgements

This work was supported by the Deutsche Forschungsgemeinschaft (DFG). We thank Eva-Maria Bornholdt and Lea Rittmeier for plasmid constructions and Gudrun Ebel and Karola Hahn for technical support.

## Declarations

### Availability of data and material

Original data are available upon request. Additional information is provided in the Supplementary Material.

### Code availability

Not applicable

### Authors’ contributions

HJS conceived the study, designed the experiments, supervised the project and wrote the manuscript. JO constructed *CAB1* variants and analyzed PanK function. MG constructed pLEUTEX plasmids and yeast strains. LS analyzed *cab1*^ts^ revertants, performed PanK assays and functional studies of h*COASY* in yeast. TU constructed yeast strains and analyzed CoA+acetyl-CoA in yeast extracts.

Ethics approval: Not applicable (no studies with human participants or animals were performed in this study).

### Consent for publication

All authors have read and approved the final manuscript.

## References

Abrie JA, González A, Strauss E, Ariño J (2012) Functional mapping of the disparate activities of the yeast moonlighting protein Hal3. Biochem J 442:357–368

Abrie JA, Molero C, Ariño J, Strauss E (2015) Complex stability and dynamic subunit interchange modulates the disparate activities of the yeast moonlighting proteins Hal3 and Vhs3. Sci Rep 5:15774

Baković J, López Martínez D, Nikolaou S, Yu BYK, Tossounian MA, Tsuchiya Y, Thrasivoulou C, Filonenko V, Gout I (2021) Regulation of the CoA biosynthetic complex assembly in mammalian cells. Int J Mol Sci 22:1131

Baković J, Yu BYK, Silva D, Chew SP, Kim S, Ahn SH, Palmer L, Aloum L, Stanzani G, Malanchuk O, Duchen MR, Singer M, Filonenko V, Lee TH, Skehel M, Gout I (2019) A key metabolic integrator, coenzyme A, modulates the activity of peroxiredoxin 5 via covalent modification. Mol Cell Biochem 461:91–102

Bergmeyer HU (1974) Methods of Enzymatic Analysis. Vol. 4, 2^nd^ edition, Academic Press, New York.

Brachmann CB, Davies A, Cost GJ, Caputo E, Li J, Hieter P, Boeke JD (1998) Designer deletion strains derived from *Saccharomyces cerevisiae* S288C: a useful set of strains and plasmids for PCR-mediated gene disruption and other applications. Yeast 14:115–132

Brohée S, Janky R, Abdel-Sater F, Vanderstocken G, André B, van Helden J (2011) Unraveling networks of co-regulated genes on the sole basis of genome sequences. Nucleic Acids Res 39:6340–6358

Calder RB, Williams RS, Ramaswamy G, Rock CO, Campbell E, Unkles SE, Kinghorn JR & Jackowski S (1999) Cloning and characterization of a eukaryotic pantothenate kinase gene *(panK)* from *Aspergillus nidulans*. J Biol Chem 274:2014–2020

Cartwright JL, Gasmi L, Spiller DG, McLennan AG (2000) The *Saccharomyces cerevisiae PCD1* gene encodes a peroxisomal nudix hydrolase active toward coenzyme A and its derivatives. J Biol Chem 275:32925–32930

Chen Y, Daviet L, Schalk M, Siewers V, Nielsen J (2013) Establishing a platform cell factory through engineering of yeast acetyl-CoA metabolism. Metab Eng 15:48–54.

Daugherty M, Polanuyer B, Farrell M, Scholle M, Lykidis A, de Crécy-Lagard V & Osterman A (2002) Complete reconstitution of the human coenzyme A biosynthetic pathway via comparative genomics. J Biol Chem 277:21431–21439

Dusi S, Valletta L, Haack TB, Tsuchiya Y, Venco P, Pasqualato S, Goffrini P, Tigano M, Demchenko N, Wieland T, Schwarzmayr T, Strom TM, Invernizzi F, Garavaglia B, Gregory A, Sanford L, Hamada J, Bettencourt C, Houlden H, Chiapparini L, Zorzi G, Kurian MA, Nardocci N, Prokisch H, Hayflick S, Gout I, Tiranti V (2014) Exome sequence reveals mutations in CoA synthase as a cause of neurodegeneration with brain iron accumulation. Am J Hum Genet 94:11–22

Galdieri L, Zhang T, Rogerson D, Lleshi R, Vancura A (2014) Protein acetylation and acetyl coenzyme a metabolism in budding yeast. Eukaryot Cell 13:1472–1483

Güldener U, Heck S, Fielder T, Beinhauer J, Hegemann JH (1996) A new efficient gene disruption cassette for repeated use in budding yeast. Nucleic Acids Res 24:2519–2524

Hayflick SJ (2014) Defective pantothenate metabolism and neurodegeneration. Biochem Soc Trans 42:1063–1068

Hong BS, Senisterra G, Rabeh WM, Vedadi M, Leonardi R, Zhang YM, Rock CO, Jackowski S, Park HW (2007) Crystal structures of human pantothenate kinases. Insights into allosteric regulation and mutations linked to a neurodegeneration disorder. J Biol Chem 282:27984–27993

Hong KQ, Fu XM, Dong SS, Xiao DG, Dong J (2019) Modulating acetate ester and higher alcohol production in *Saccharomyces cerevisiae* through the cofactor engineering. J Ind Microbiol Biotechnol 46:1003–1011

Iuso A, Wiersma M, Schüller HJ, Pode-Shakked B, Marek-Yagel D, Grigat M, Schwarzmayr T, Berutti R, Alhaddad B, Kanon B, Grzeschik NA, Okun JG, Perles Z, Salem Y, Barel O, Vardi A, Rubinshtein M, Tirosh T, Dubnov-Raz G, Messias AC, Terrile C, Barshack I, Volkov A, Avivi C, Eyal E, Mastantuono E, Kumbar M, Abudi S, Braunisch M, Strom TM, Meitinger T, Hoffmann GF, Prokisch H, Haack TB, Brundel BJJM, Haas D, Sibon OCM, Anikster Y. Mutations in PPCS, encoding Phosphopantothenoylcysteine Synthetase, cause autosomal-recessive dilated cardiomyopathy. Am J Hum Genet 102:1018–1030

Leonardi R, Chohnan S, Zhang YM, Virga KG, Lee RE, Rock CO, Jackowski S (2005a) A pantothenate kinase from *Staphylococcus aureus* refractory to feedback regulation by coenzyme A. J Biol Chem 280:3314–3322

Leonardi R, Zhang YM, Rock CO & Jackowski S (2005b) Coenzyme A: back in action. Prog Lipid Res 44:125–153

Lian J, Si T, Nair NU, Zhao H (2014) Design and construction of acetyl-CoA overproducing *Saccharomyces cerevisiae* strains. Metab Eng 24:139–149

Luo X, Reiter MA, d’Espaux L, Wong J, Denby CM, Lechner A, Zhang Y, Grzybowski AT, Harth S, Lin W, Lee H, Yu C, Shin J, Deng K, Benites VT, Wang G, Baidoo EEK, Chen Y, Dev I, Petzold CJ, Keasling JD (2019) Complete biosynthesis of cannabinoids and their unnatural analogues in yeast. Nature 567:123–126

Meadows AL, Hawkins KM, Tsegaye Y, Antipov E, Kim Y, Raetz L, Dahl RH, Tai A, Mahatdejkul-Meadows T, Xu L, Zhao L, Dasika MS, Murarka A, Lenihan J, Eng D, Leng JS, Liu CL, Wenger JW, Jiang H, Chao L, Westfall P, Lai J, Ganesan S, Jackson P, Mans R, Platt D, Reeves CD, Saija PR, Wichmann G, Holmes VF, Benjamin K, Hill PW, Gardner TS, Tsong AE (2016) Rewriting yeast central carbon metabolism for industrial isoprenoid production. Nature 537:694–697

Milke L, Marienhagen J (2020) Engineering intracellular malonyl-CoA availability in microbial hosts and its impact on polyketide and fatty acid synthesis. Appl Microbiol Biotechnol 104:6057–6065

Mumberg D, Müller R & Funk M (1994) Regulatable promoters of *Saccharomyces cerevisiae*: comparison of transcriptional activity and their use for heterologous expression. Nucleic Acids Res 22:5767–5768

Naquet P, Kerr EW, Vickers SD, Leonardi R (2020) Regulation of coenzyme A levels by degradation: the ‘Ins and Outs’. Prog Lipid Res 78:101028

Olzhausen J, Moritz T, Neetz T, Schüller HJ (2013) Molecular characterization of the heteromeric coenzyme A-synthesizing protein complex (CoA-SPC) in the yeast *Saccharomyces cerevisiae*. FEMS Yeast Res 13:565–573

Olzhausen J, Schübbe S & Schüller HJ (2009) Genetic analysis of coenzyme A biosynthesis in the yeast *Saccharomyces cerevisiae*: identification of a conditional mutation in the pantothenate kinase gene *CAB1*. Curr Genet 55:163–173

Paddon CJ, Westfall PJ, Pitera DJ, Benjamin K, Fisher K, McPhee D, Leavell MD, Tai A, Main A, Eng D, Polichuk DR, Teoh KH, Reed DW, Treynor T, Lenihan J, Fleck M, Bajad S, Dang G, Dengrove D, Diola D, Dorin G, Ellens KW, Fickes S, Galazzo J, Gaucher SP, Geistlinger T, Henry R, Hepp M, Horning T, Iqbal T, Jiang H, Kizer L, Lieu B, Melis D, Moss N, Regentin R, Secrest S, Tsuruta H, Vazquez R, Westblade LF, Xu L, Yu M, Zhang Y, Zhao L, Lievense J, Covello PS, Keasling JD, Reiling KK, Renninger NS, Newman JD (2013) High-level semi-synthetic production of the potent antimalarial artemisinin. Nature 496:528–532

Pietrocola F, Galluzzi L, Bravo-San Pedro JM, Madeo F, Kroemer G (2015) Acetyl coenzyme A: a central metabolite and second messenger. Cell Metab 21:805–821

Rock CO, Calder RB, Karim MA, Jackowski S (2000) Pantothenate kinase regulation of the intracellular concentration of coenzyme A. J Biol Chem 275:1377–1383

Rock CO, Park HW, Jackowski S (2003) Role of feedback regulation of pantothenate kinase (CoaA) in control of coenzyme A levels in *Escherichia coli*. J Bacteriol 185:3410–3415

Rothstein R (1991) Targeting, disruption, replacement, and allele rescue: integrative DNA transformation in yeast. Methods Enzymol 194:281–301

Ruiz A, González A, Muñoz I, Serrano R, Abrie JA, Strauss E & Ariño J (2009) Moonlighting proteins Hal3 and Vhs3 form a heteromeric PPCDC with Ykl088w in yeast CoA biosynthesis. Nat Chem Biol 5:920–928

Sandoval CM, Ayson M, Moss N, Lieu B, Jackson P, Gaucher SP, Horning T, Dahl RH, Denery JR, Abbott DA, Meadows AL (2014) Use of pantothenate as a metabolic switch increases the genetic stability of farnesene producing *Saccharomyces cerevisiae*. Metab Eng 25:215–226

Schadeweg V, Boles E (2016) n-Butanol production in *Saccharomyces cerevisiae* is limited by the availability of coenzyme A and cytosolic acetyl-CoA. Biotechnol Biofuels 9:44

Shi L, Tu BP (2013) Acetyl-CoA induces transcription of the key G1 cyclin *CLN3* to promote entry into the cell division cycle in *Saccharomyces cerevisiae*. Proc Natl Acad Sci USA 110:7318–7323

Shi L, Tu BP (2015) Acetyl-CoA and the regulation of metabolism: mechanisms and consequences. Curr Opin Cell Biol 33:125–131

Sibon OCM, Strauss E (2016) Coenzyme A: to make it or uptake it? Nat Rev Mol Cell Biol 17:605–606

Sikorski RS, Boeke JD (1991) In vitro mutagenesis and plasmid shuffling: from cloned gene to mutant yeast. Methods Enzymol 194:302–318

Sikorski RS, Hieter P (1989) A system of shuttle vectors and yeast host strains designed for efficient manipulation of DNA in *Saccharomyces cerevisiae*. Genetics 122:19–27

Song WJ, Jackowski S (1992) Cloning, sequencing, and expression of the pantothenate kinase *(coaA)* gene of *Escherichia coli*. J Bacteriol 174:6411–6417

Spry C, Kirk K, Saliba KJ (2008) Coenzyme A biosynthesis: an antimicrobial drug target. FEMS Microbiol Rev 32:56–106

Srinivasan B, Baratashvili M, van der Zwaag M, Kanon B, Colombelli C, Lambrechts RA, Schaap O, Nollen EA, Podgoršek A, Kosec G, Petković H, Hayflick S, Tiranti V, Reijngoud DJ, Grzeschik NA, Sibon OC (2015) Extracellular 4’-phosphopantetheine is a source for intracellular coenzyme A synthesis. Nat Chem Biol 11:784–792

Stolz J, Sauer N (1999) The fenpropimorph resistance gene *FEN2* from *Saccharomyces cerevisiae* encodes a plasma membrane H^+^-pantothenate symporter. J Biol Chem 274:18747–18752

Stolz J, Caspari T, Carr AM, Sauer N (2004) Cell division defects of *Schizosaccharomyces pombe liz1*^-^ mutants are caused by defects in pantothenate uptake. Eukaryot Cell 3:406–412

Szczebara FM, Chandelier C, Villeret C, Masurel A, Bourot S, Duport C, Blanchard S, Groisillier A, Testet E, Costaglioli P, Cauet G, Degryse E, Balbuena D, Winter J, Achstetter T, Spagnoli R, Pompon D, Dumas B (2003) Total biosynthesis of hydrocortisone from a simple carbon source in yeast. Nat Biotechnol 21:143–149

Theodoulou FL, Sibon OC, Jackowski S, Gout I (2014) Coenzyme A and its derivatives: renaissance of a textbook classic. Biochem Soc Trans 42:1025–1032

Tippmann S, Scalcinati G, Siewers V, Nielsen J (2016 Production of farnesene and santalene by *Saccharomyces cerevisiae* using fed-batch cultivations with *RQ*-controlled feed. Biotechnol Bioeng 113:72–81

Vallari DS, Jackowski S & Rock CO (1987) Regulation of pantothenate kinase by coenzyme A and its thioesters. J Biol Chem 262:2468–2471

White WH, Gunyuzlu PL & Toyn JH (2001) *Saccharomyces cerevisiae* is capable of *de novo* pantothenic acid biosynthesis involving a novel pathway of β-alanine production from spermine. J Biol Chem 276:10794–10800

Zhang YM, Rock CO, Jackowski S (2005) Feedback regulation of murine pantothenate kinase 3 by coenzyme A and coenzyme A thioesters. J Biol Chem 280:32594–32601

Zhyvoloup A, Nemazanyy I, Babich A, Panasyuk G, Pobigailo N, Vudmaska M, Naidenov V, Kukharenko O, Palchevskii S, Savinska L, Ovcharenko G, Verdier F, Valovka T, Fenton T, Rebholz H, Wang ML, Shepherd P, Matsuka G, Filonenko V, Gout IT (2002) Molecular cloning of CoA Synthase: The missing link in CoA biosynthesis. J Biol Chem 277:22107–22110

Zhyvoloup A, Nemazanyy I, Panasyuk G, Valovka T, Fenton T, Rebholz H, Wang ML, Foxon R, Lyzogubov V, Usenko V, Kyyamova R, Gorbenko O, Matsuka G, Filonenko V, Gout IT (2003) Subcellular localization and regulation of coenzyme A synthase. J Biol Chem 278:50316–50321

